# Revealing A-T and G-C Hoogsteen base pairs in stressed protein-bound duplex DNA

**DOI:** 10.1101/2021.06.05.447203

**Authors:** Honglue Shi, Issac J. Kimsey, Hsuan-Fu Liu, Uyen Pham, Maria A. Schumacher, Hashim M. Al-Hashimi

## Abstract

Watson-Crick base pairs (bps) are the fundamental unit of genetic information and the building blocks of the DNA double helix. However, A-T and G-C can also form alternative ‘Hoogsteen’ bps, expanding the functional complexity of DNA. We developed ‘Hoog-finder’, which uses structural fingerprints to rapidly screen Hoogsteen bps, which may have been mismodeled as Watson-Crick in crystal structures of protein-DNA complexes. We uncovered seventeen Hoogsteen bps, seven of which were in complex with six proteins never before shown to bind Hoogsteen bps. The Hoogsteen bps occur near mismatches, nicks, and lesions and some appear to participate in recognition and damage repair. Our results suggest a potentially broad role for Hoogsteen bps in stressed regions of the genome and call for a community-wide effort to identify these bps in current and future crystal structures of DNA and its complexes.

## Introduction

One of the cornerstones of molecular biology is that A pairs with T and G with C to form Watson-Crick base pairs (bps) (Fig. 1a). However, soon after the discovery of the DNA double helix, it was shown that A-T and G-C could also pair in an alternative conformation known as the ‘Hoogsteen’ bp^1, 2^ (Fig. 1a). A Hoogsteen bp can be obtained by flipping the purine base in a Watson-Crick bp from the *anti* to *syn* conformation and then forming a unique set of hydrogen bonds (H-bonds) with the partner pyrimidine requiring protonation of cytosine-N3 (Fig. 1a). Relative to Watson-Crick bps, Hoogsteen pairing requires that the two bases also come into closer proximity by ∼2.0-2.5 Å. This has been shown to locally constrict the helical diameter and to cause kinking of the DNA double helix toward the major groove by ∼10°^3^.

**Fig. 1.**
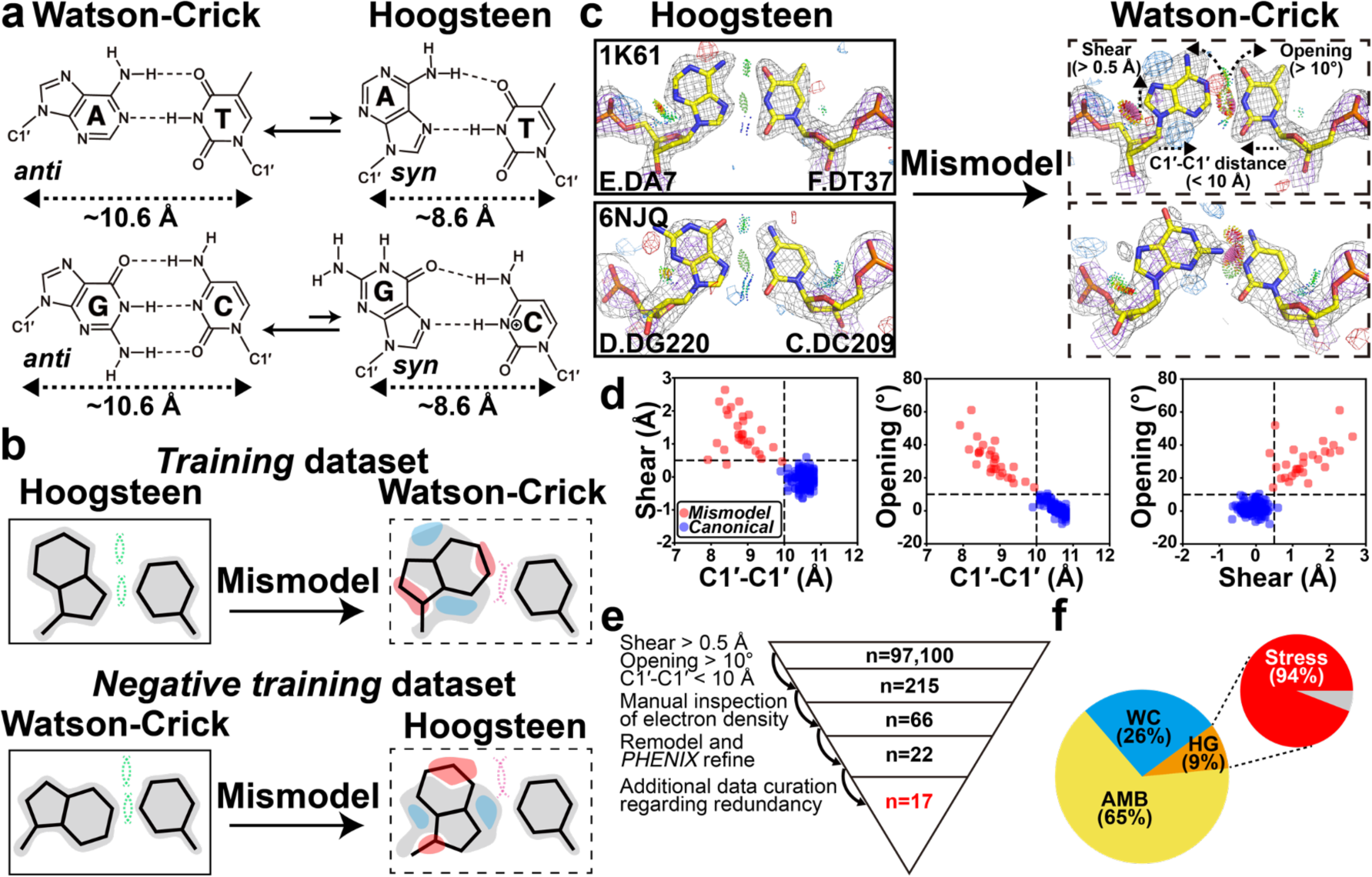
Hoog-finder to rapidly identify putative Hoogsteen bps in crystal structures of protein-DNA complexes. **(a)** Dynamic equilibrium between Watson-Crick and Hoogsteen bps. **(b)** Generating the training and negative training datasets. 2mF_o_-DF_c_ electron density maps (contoured at ∼1 σ) are shown in gray, whereas red and blue regions represent mF_o_-DF_c_ difference electron density maps contoured at around +3σ and −3σ, respectively. Steric clashes and H-bonds between the two bases are denoted using a pink and a green dashed line, respectively. **(c)** Representative 2mF_o_-DF_c_ and mF_o_-DF_c_ electron density maps for original Hoogsteen (left, solid boxes) and the corresponding mismodeled Watson-Crick models (right, dashed boxes) highlighting the unique structural fingerprints of mismodeled Watson-Crick bps. Gray and purple meshed regions represent 2mF_o_-DF_c_ densities at 1.0σ and 3.0σ, respectively, while blue and red meshed regions are mF_o_-DF_c_ difference densities contoured at 3.0σ and −3.0σ, respectively. Also shown is the stereochemistry assessed by *MolProbity* (Methods). All bp structures and electron densities in the *training* dataset are provided in Extended Data Fig. 1. **(d)** 2D scatter plot comparing C1′-C1′ distance, shear, and opening for Hoogsteen bps when is mismodeled as Watson-Crick (red, n=28) and the canonical Watson-Crick dataset^16^ (blue, n=149). The three structural criteria are denoted as the dashed line. **(e)** Workflow used to identify putative Hoogsteen bps mismodeled as Watson-Crick. **(f)** Percentage distribution of bps identified to be Hoogsteen (HG, orange), Watson-Crick (WC, skyblue) and ambiguous bps (AMB, yellow). Data shown for non-redundant bps following data curation (Methods). Also shown is the percentage of Hoogsteen bps found in stressed regions of DNA.

Following their initial discovery, Hoogsteen bps were observed in a handful of crystal structures of protein-DNA complexes and shown to participate in DNA shape recognition^4–7^. An early example was the crystal structure (PDB: 1IHF) of duplex DNA in complex with the integration host factor (IHF) protein^4^. The structure included an unusual A(*anti*)-T Hoogsteen bp in which the adenine base was in the *anti* rather than *syn* conformation. The bp was located immediately adjacent to a nick used to aid crystallization^4^.

More conventional A(*syn*)-T and G(*syn*)-C^+^ Hoogsteen bps in which the purine base is in the *syn* conformation were subsequently reported in crystal structures of intact DNA duplexes in complex with transcription factors, including the TATA box-binding protein (TBP)^5^ (PDB: 1QN3, 6NJQ), MAT*α*2 homeodomain^6^ (PDB: 1K61), and the DNA binding domain of the p53 tumor suppressor protein^7^ (PDB: 3KZ8). Beyond transcription factors, crystallographic and biochemical studies also revealed Hoogsteen bps in the active sites of specialized polymerases including human polymerase ι^8, 9^ (PDB: 1TN3, 2ALZ) and *Sulfolobus solfataricus* polymerase Dpo4^10, 11^ (PDB: 1RYS, 1S0M), in which they were proposed to be involved in mediating the bypass of DNA damage during replication. These crystal structures together with structures of certain DNA-drug complexes^12^ established Hoogsteen bps as an alternative to Watson-Crick imparting unique characteristics to the DNA.

NMR studies later revealed Hoogsteen bps are ubiquitous in DNA duplexes. Across a wide variety of sequence and positional contexts, A-T and G-C Watson-Crick bps were shown to exist in dynamic equilibrium with their Hoogsteen counterparts^13^ (Fig. 1a). The population (∼0.1%-1.0%) of the minor Hoogsteen conformation exceeds that of other conformational states commonly stabilized by proteins such as the base open conformation^14^ by more than two orders of magnitude. Since Hoogsteen bps can also occur in any sequence context^15^, it is surprising that they have not been more extensively observed in crystal structures of DNA, particularly in protein-DNA complexes, in which the DNA structure is often highly distorted and conformationally stressed. Indeed, Hoogsteen bps appear to favor stressed regions in which the helix is unwound and/or kinked toward the major groove as well as at terminal ends of the DNA^16, 17^ and in which neighboring bps are partially melted^3^.

Prior crystallographic studies have underscored the difficulty distinguishing Watson-Crick from Hoogsteen bps especially when the electron density is of moderate or low quality^7, 8, 18–20^. Because a Watson-Crick bp is generally assumed initially unless there is other data to indicate otherwise, or the structure is at high resolution and reveals a clear non-Watson-Crick conformation, some of the Watson-Crick bps in current crystal structures of DNA in the Protein Data Bank (PDB)^21^ could be ambiguous. Some might even be better modeled as Hoogsteen bps.

Re-analyzing the electron density for some ∼100,000 DNA bps bound to proteins in the PDB to assess the degree to which the data supports the Watson-Crick versus a Hoogsteen model is laborious and impractical. To help streamline this analysis, a recent study^20^ developed an automated approach, which uses differences in electron density expected for Watson-Crick versus Hoogsteen bp models as fingerprints to identify Hoogsteen bps mismodeled as Watson-Crick. This work identified eight Hoogsteen bps mismodeled as Watson-Crick at terminal ends of DNA sites and in structures of DNA in complex with the polymerase Dpo4 which had previously been shown to bind DNA with Hoogsteen bps at certain positions^10, 11, 22, 23^.

Here, we developed an alternative structure-guided approach termed ‘Hoog-finder’ to rapidly screen for Hoogsteen bps that may have been mismodeled as Watson-Crick in crystal structures of protein-DNA complexes. Using Hoog-finder, we uncovered 17 bps that better satisfy the electron density and also result in improved stereochemistry when modeled as Hoogsteen relative to Watson-Crick. Seven of these Hoogsteen bps were observed in DNA in complexes of six proteins never before shown to bind DNA in a Hoogsteen conformation. Interestingly, almost all of the newly uncovered Hoogsteen bps were adjacent to mismatches, lesions, nicks, and terminal ends, and some of them appear to play roles in DNA recognition and damage repair. In addition, more than half of the ∼200 bps examined had ambiguous electron density. Among these, 21 bps had slightly better fits to the electron density and/or resulted in improved stereochemistry when modeled as Hoogsteen relative to Watson-Crick. Thus, our results point to potentially broader roles for Hoogsteen bps than currently appreciated, particularly in stressed regions of the genome, and call for a community-wide effort to identify these bps in current and future crystal structures of DNA.

## Results

### Structural fingerprints of Hoogsteen bps mismodeled as Watson-Crick

We hypothesized that mismodeling a Watson-Crick bp into electron density belonging to a Hoogsteen bp could result in a distorted Watson-Crick geometry deviating from the canonical Watson-Crick conformation (Fig. 1b-d). These geometrical distortions could then be used as ‘structural fingerprints’ to screen the PDB for Hoogsteen bps that had been mismodeled as Watson-Crick before examining the electron density, which is laborious and time-consuming.

To examine whether or not Hoogsteen bps mismodeled as Watson-Crick have unique geometrical distortions, we built a *training* dataset (Supplementary Table 1) of previously reported A(*syn*)-T and G(*syn*)-C^+^ Hoogsteen bps^16^ with available structure factors. The dataset comprised 22 A(*syn*)-T and six G(*syn*)-C^+^ Hoogsteen bps from 23 crystal structures of duplex DNA, 22 were DNA-protein complexes and one was a naked DNA duplex (Supplementary Table 1).

For each Hoogsteen bp in the *training* dataset, we generated an *omit* map by removing the *syn* purine. We also deliberately mismodeled the Watson-Crick bp by introducing a purine residue in the *anti* conformation. The resulting structure was refined using *PHENIX*^24, 25^ to generate coordinates and electron density maps for the structure with a mismodeled Watson-Crick bp (Fig. 1c, Extended Data Fig. 1 and Methods). Except for two bps, which had ambiguous electron density, the Hoogsteen bps showed better agreement with the electron density and better stereochemistry when assessed by *MolProbity*^26^ compared to the Watson-Crick bps (Fig. 1c, Extended Data Fig. 1, Supplementary Note 1 and Methods). However, the extent of improvement varied from case to case, in agreement with the original publications.

We then compared the geometrical features of the mismodeled Watson-Crick bps with those of canonical Watson-Crick bps. The canonical Watson-Crick geometry was defined based on n=149 bps obtained from a prior survey^16^ (Methods) with well-defined density satisfying the Watson-Crick geometry. The geometrical features analyzed included backbone torsion angles, sugar pucker, bp parameters, C1′-C1′ inter-nucleotide distance, as well as major and minor groove widths (Extended Data Fig. 2a-c).

**Fig. 2.**
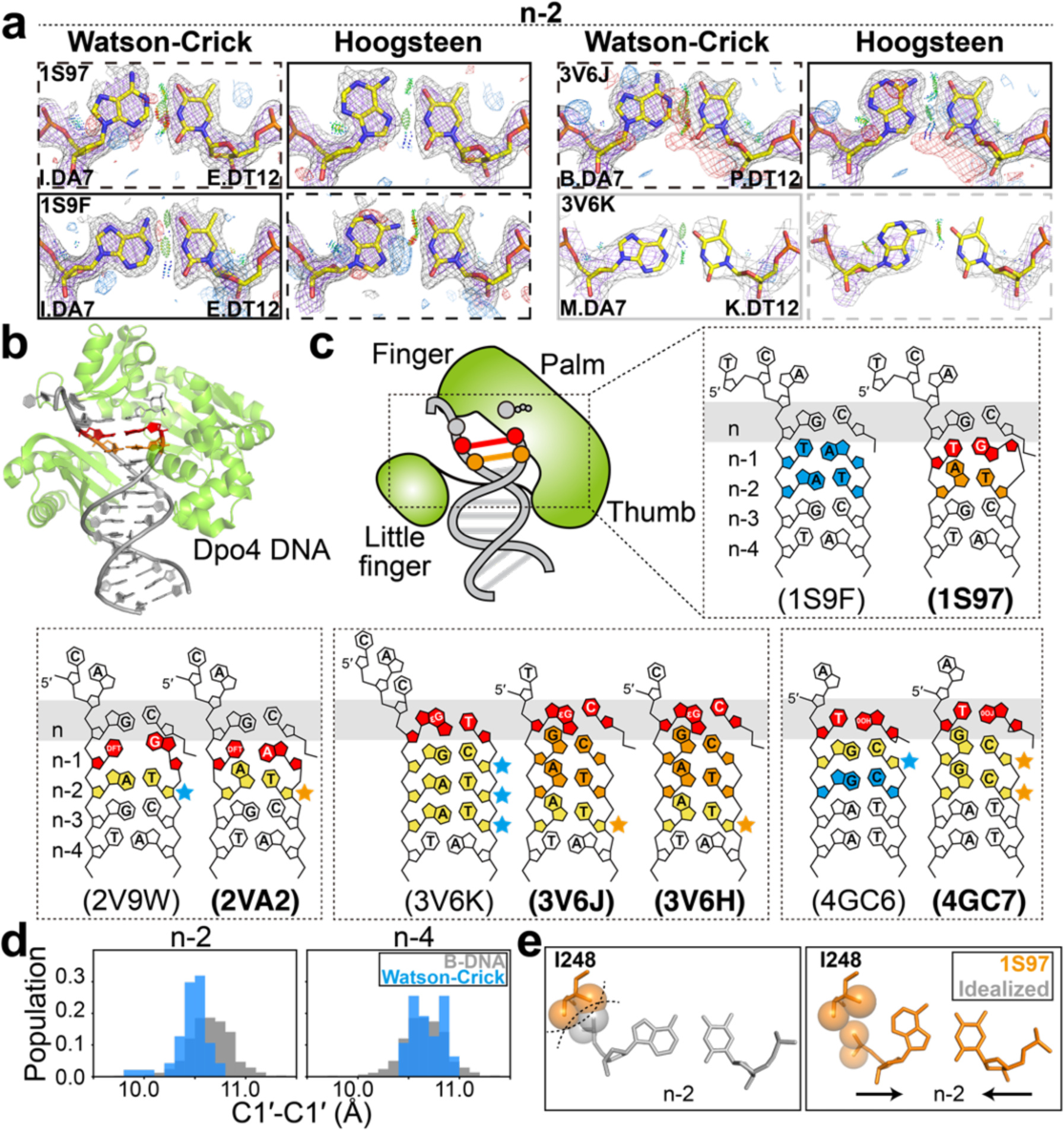
Hoogsteen base pairs in Dpo4. **(a)** Comparison of 2mF_o_-DF_c_ and mF_o_-DF_c_ electron density maps calculated with models containing the original Watson-Crick (left) and Hoogsteen models (right). Electron density meshes and stereochemistry are as described in Fig. 1c. The favored and less favored models are indicated using solid and dashed boxes, respectively. The boxes are in gray for ambiguous bps. A complete set of data is provided in Extended Data Fig. 6-8. **(b)** 3D structures of the protein-DNA complex showing the Hoogsteen bps. **(c)** Schematic showing the DNA (bolded PDB ID) containing Hoogsteen bps (in orange) and ambiguous Hoogsteen bps (in yellow next to orange stars). The corresponding structures (unbolded PDB ID) containing Watson-Crick bps (in skyblue) and ambiguous Watson-Crick bps (in yellow next to skyblue stars) are also shown inside the same dashed box. The lesions and mismatches were highlighted in red. **(d)** Distribution of C1′-C1′ distance between bases in bps at positions *n*-2 and *n*-4 in the Dpo4 DNA without lesions or mismatches (see PDB ID in Supplementary Table 8) (skyblue), showing constriction at *n*=-2 compared to distribution of Watson-Crick bps in B-DNA (in gray) from Afek *et al.*^30^. **(e)** Close up of the Hoogsteen bp in the active site of Dpo4, which are more compressed than idealized B-form DNA, thus avoiding potential steric clash between the DNA backbone at *n*-2 and Ile248.

For most structural parameters, including backbone torsion angles, sugar pucker, and groove widths, we did not observe a clear distinction between the mismodeled and canonical Watson-Crick bps (Extended Data Fig. 2d-f). However, for all the mismodeled Watson-Crick bps, the C1′-C1′ distance was consistently reduced by more than 0.6 Å (from ∼10.6 Å to <10.0 Å) relative to the canonical Watson-Crick geometry (Fig. 1c-d and Extended Data Fig. 2d). Constriction of the C1′-C1′ distance by ∼2.0-2.5 Å has been shown to be one of the most distinguishing structural^3, 27^ as well as functional^28–30^ characteristics of the Hoogsteen bps relative to Watson-Crick, and it is not surprising that modelling Watson-Crick bps into density belonging to Hoogsteen bps would result in a constriction (Fig. 1a). In addition, the purine base was also consistently displaced towards the major groove (shear >0.5 Å) and adopted a more open conformation (opening >10°) relative to a canonical Watson-Crick bp (Fig. 1d and Extended Data Fig. 2d). These deviations likely accommodate constriction of the C1′-C1′ distance, without them, the two bases would sterically clash.

Conversely, using a *negative training* dataset (n=10) of Watson-Crick bps, we also asked whether there were ‘structural fingerprints’, which could be used to identify cases in which a Watson-Crick bp was mismodeled as Hoogsteen (Fig. 1b). Indeed, we found that such mismodeled Hoogsteen bps have C1′-C1′ distances exceeding 10.0 Å, with the *syn* purine base being substantially displaced towards the minor groove resulting in loss of the H-bond between the purine-N7 and pyrimidine-N3 and oftentimes resulting in steric clashes between purine-N6/O6 and pyrimidine-N4/O4/N3 (Extended Data Fig. 3, Supplementary Table 2 and Methods).

**Fig. 3.**
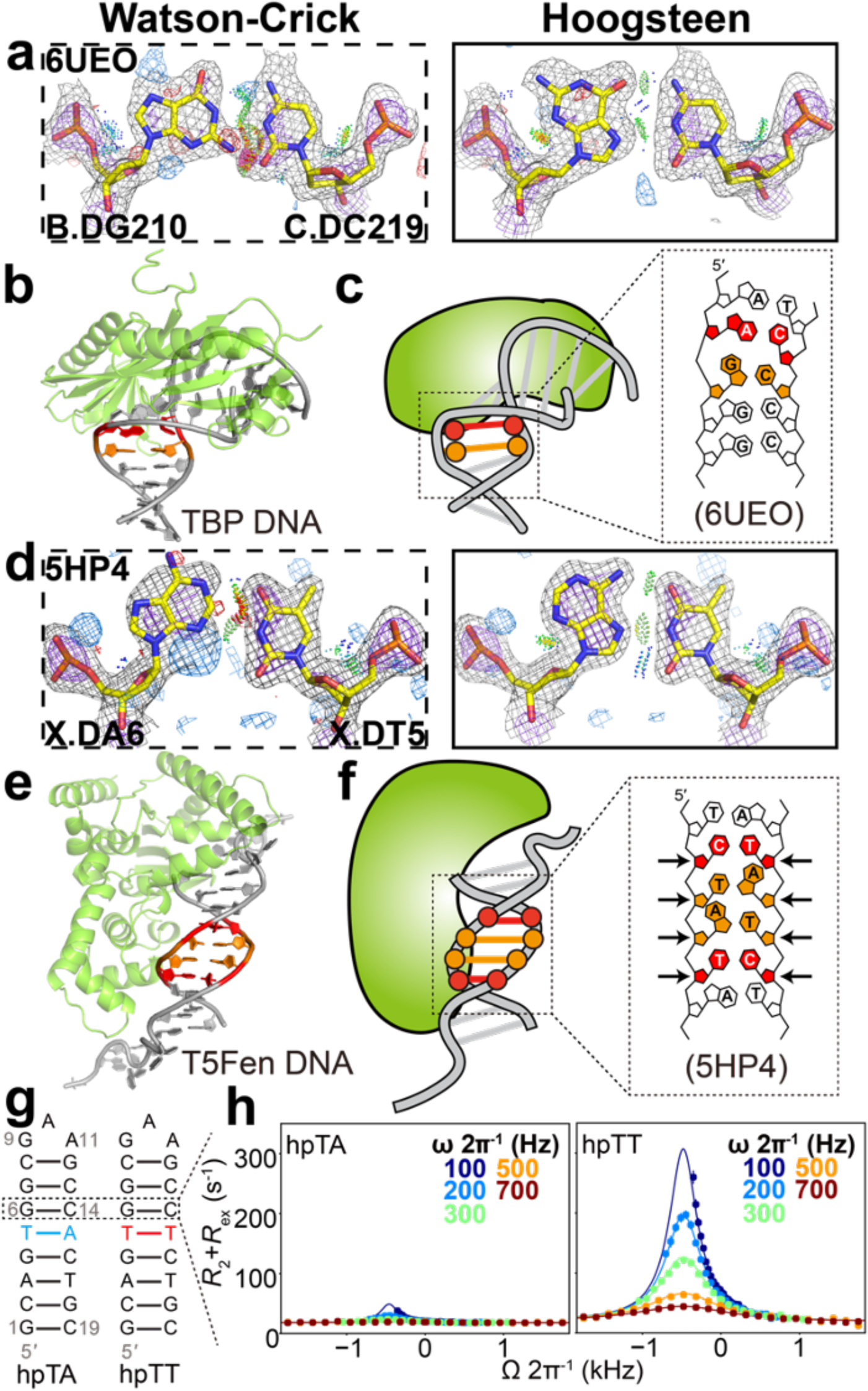
Hoogsteen base pairs next to mismatches. **(a,d)** Comparison of 2mF_o_-DF_c_ and mF_o_-DF_c_ electron density maps for the Watson-Crick (left) and the corresponding Hoogsteen models (right) for **(a)** the G-C bp next to an A-C mismatch in TBP, and **(d)** the A-T bp next to a C-T mismatch in T5-flap endonuclease. Note that the G-C bp in **(a)** was modeled as a Hoogsteen bp in PDB 6UEO. Electron density meshes and stereochemistry are as described in Fig. 1c and the box scheme is as described in Fig. 2a. **(b,e)** 3D structures of the protein-DNA complex showing the Hoogsteen bps for **(b)** TBP and **(e)** T5-flap endonuclease. **(c,f)** Schematic showing the DNA containing Hoogsteen bps (in orange), as well as the mismatches (in red) for **(c)** TBP and **(f)** T5-flap endonuclease. **(g)** Hairpin DNA with and without mismatch used in NMR measurements. Hoogsteen populations were measured at G6-C14 bp. **(h)** NMR off-resonance *R*_1ρ_ profiles of G6-C8. Spin-lock powers are color coded. Error bars were estimated using a Monte-Carlo scheme and are smaller than data points (Methods).

Based on these results, we developed ‘Hoog-finder’, a structure-based approach to rapidly identify Hoogsteen bps which may have been mismodeled as Watson-Crick. Such bps could be identified if they satisfied all three ‘positive structural fingerprints’ (C1′-C1′ distance <10 Å, shear >0.5 Å, and opening >10°) while also not satisfying the ‘negative structural fingerprints’ after being remodeled as Hoogsteen bps. As an initial test, Hoog-finder identified seven of eight Hoogsteen bps that were mismodeled as Watson-Crick and identified in a prior analysis^20^, with the one exception only satisfying two of the positive structural fingerprints.

### Structure-based approach for identifying Hoogsteen bps mismodeled as Watson-Crick

We used Hoog-finder to screen 97,100 Watson-Crick bps in a *Parent* dataset (n=97,100) representing 4,002 crystal structures of all DNA-protein complexes in the PDB as of Aug 29^th^, 2020 with resolution better than 3.5 Å. Hoog-finder identified 215 Watson-Crick bps in 173 crystal structures (Fig. 1e and Methods). Pseudo-palindromic DNA sites, which displayed possible statistical disorder^20^, were not included in the analysis (Methods). The electron density for each of these bps was then analyzed manually.

Of the Watson-Crick 215 bps examined, 58 showed good agreement with electron density and favorable stereochemistry as assessed using *MolProbity* (Extended Data Fig. 4a and Supplementary Tables 3, 4). These bps were annotated as ‘Watson-Crick’. These Watson-Crick bps were slightly distorted with geometrical features falling at the edge of the cutoff for all three structural criteria (Extended Data Fig. 5). For 91 bps, the electron density around the bp in question was too weak to evaluate the Hoogsteen or Watson-Crick model (Extended Data Fig. 4b and Supplementary Tables 3, 4). These bps were annotated as ‘ambiguous’.

**Fig. 4.**
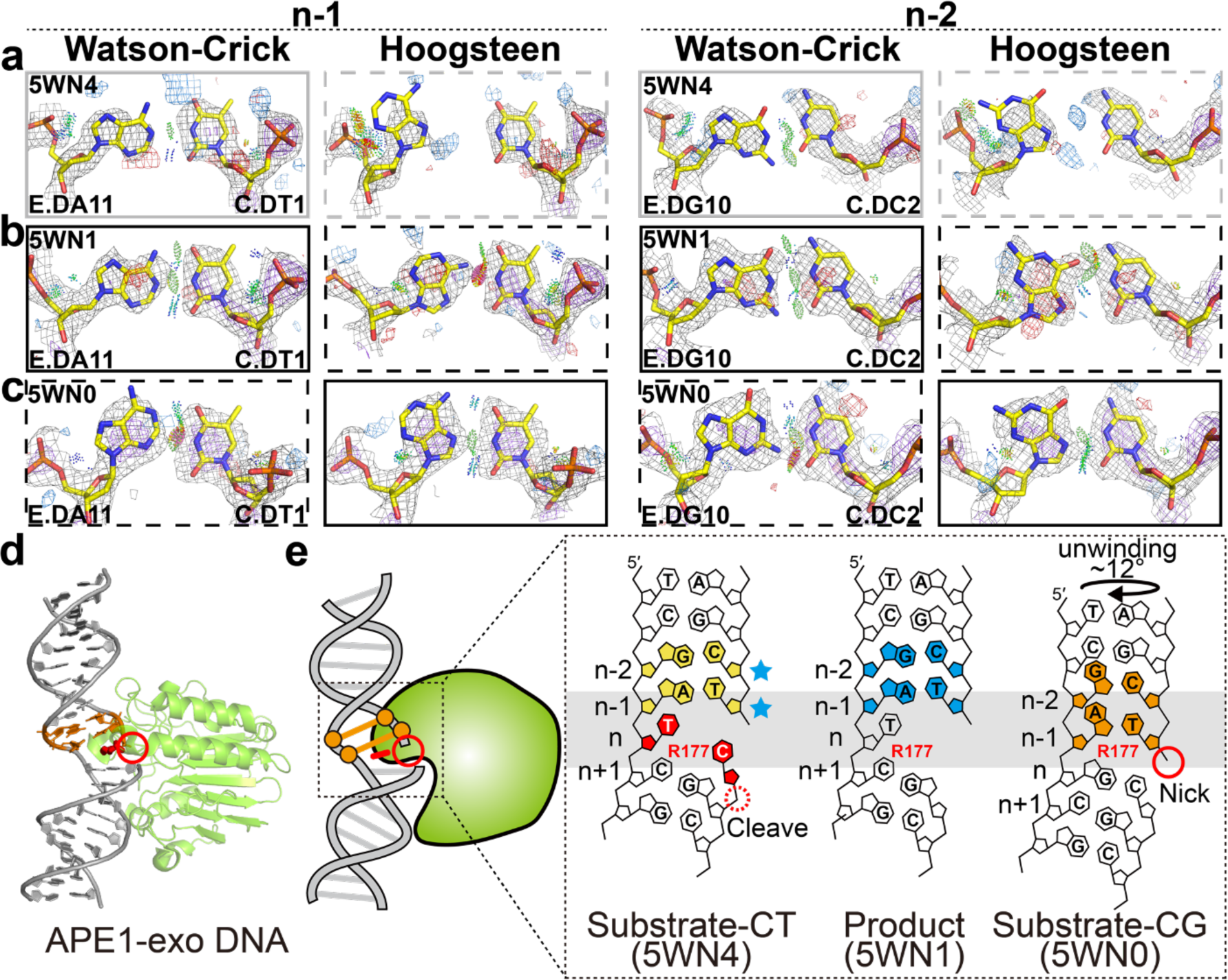
Hoogsteen base pairs in APE1-exo. **(a-c)** Comparison of 2mF_o_-DF_c_ and mF_o_-DF_c_ electron density maps for the original Watson-Crick (left) and corresponding Hoogsteen models (right) for the A-T bp at position *n*-1 and G-C bp at position *n*-2 **(a,b)** with C-T mismatch at position *n* in **(a)** a substrate complex (PDB: 5WN4) and **(b)** a product complex (PDB: 5WN1), and **(c)** with C-G match at position *n* in a substrate complex (PDB: 5WN0). Electron density meshes and stereochemistry are as described in Fig. 1c and the box scheme is as described in Fig. 2a. **(d)** 3D structures of the protein-DNA complex showing the Hoogsteen bps. **(e)** Schematic showing the DNA containing Hoogsteen bps (in orange), Watson-Crick bps (in skyblue) and ambiguous Watson-Crick bps (in yellow next to skyblue stars), as well as the R177 and mismatches (in red). Also shown is the nick (red solid circle) and cleavage site (red dashed circle).

**Fig. 5.**
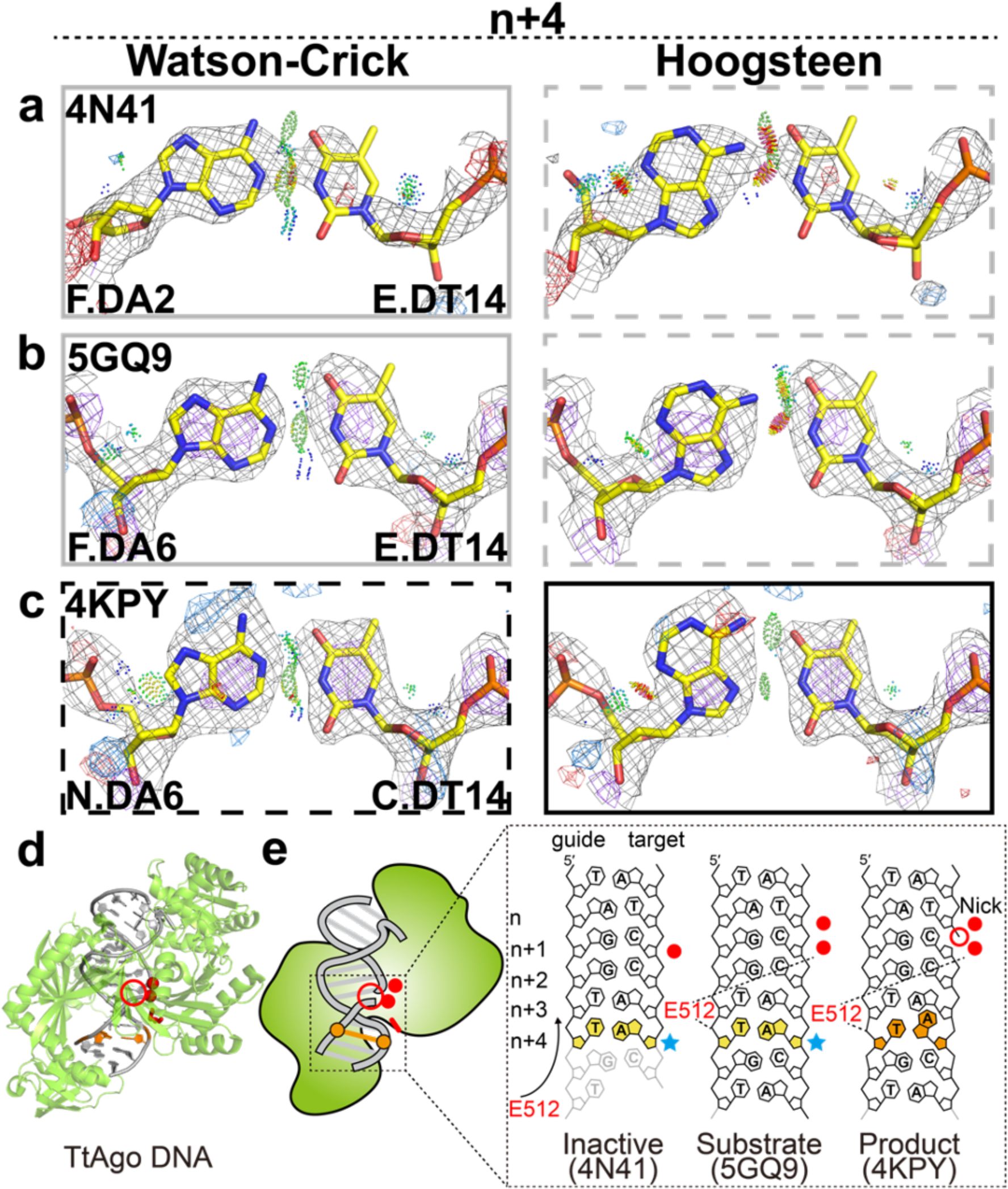
Hoogsteen base pairs in TtAgo. **(a-c)** Comparison of 2mF_o_-DF_c_ and mF_o_-DF_c_ electron density maps for the original Watson-Crick (left) and the corresponding Hoogsteen models for the A-T bp at position *n*+4 in **(a)** an inactive substrate complex with a 15-mer target DNA, **(b)** an active substrate complex with a 16-mer target DNA, **(c)** a product complex with a 19-mer target DNA with a nick between positions *n* and *n*+1. Electron density meshes and stereochemistry are as described in Fig. 1c and the box scheme is as described in Fig. 2a. A complete set of data for other bps and structures is provided in Extended Data Fig. 7. **(d)** 3D structures of the protein-DNA complex showing the Hoogsteen bps. **(e)** Schematic showing the DNA containing Hoogsteen bps (in orange), ambiguous Watson-Crick bps (in yellow next to skyblue stars), as well as the E512 (in red). Also shown is the nick (red unfilled circle) and metal ions (red filled circle). Disordered DNA regions were denoted with transparency.

The remaining 66 bps were refined using *PHENIX* to compare the Hoogsteen containing model versus the model containing a Watson-Crick bp (Fig. 1e). The resolutions of these structures ranged from 1.7 Å to 3.2 Å but most were better than 2.5 Å (Supplementary Table 5). However, while the electron densities surrounding the bps were, as expected, generally better in the high-resolution structures, the quality of the electron density varied locally around each bp and thus had to be analyzed in a case-by-case manner.

For each bp, we first generated an *omit* map by removing the *anti* purine residue. We then introduced a *syn* purine residue and refined the structure using *PHENIX* (Methods). We then assessed the agreement with the electron density maps (in particular the *omit* maps) between the refined Hoogsteen and original Watson-Crick model as well as the stereochemistry of the two bps. Bps showing much better agreement with the electron density in either Watson-Crick or Hoogsteen conformations were annotated as ‘Watson-Crick’ and ‘Hoogsteen’, respectively. In general, these bps showed better stereochemistry with the model that best fits the electron density. Bps showing a slight preference with the electron density and/or improved stereochemistry either due to lower number of steric clashes or more favorable H-bonding were labeled as ‘ambiguous Watson-Crick’ and ‘ambiguous Hoogsteen’. If no preference was observed, the bp was again labeled ‘ambiguous’. A list with all annotated bps is provided in Supplementary Table 4.

Interestingly, among these 66 bps examined, 22 showed better agreement with the electron density and stereochemistry when modeled as Hoogsteen relative to Watson-Crick (see Figs. 1e, 2a, 3d, 4c, 5c, 6b, 7a-b and Extended Data Fig. 6). As in the training dataset (Extended Data Fig. 1), the improved agreement with the electron density varied from case to case, in some cases the improvement was very substantial (e.g. PDB 5A0W in Fig. 6b) whereas in other cases the Hoogsteen was clearly the better model but the difference relative to Watson-Crick was not as strong (e.g. PDB 5WN0 in Fig. 4c). Except for nine terminal Hoogsteen bps, all of which formed crystal contacts, there were no crystal contacts observed with the remaining 13 non-terminal Hoogsteen bps that were identified. The other 44 bps were ambiguous (Extended Data Fig. 4c and Supplementary Note 2), with 23 showing slightly better agreement with Hoogsteen (‘ambiguous Hoogsteen’) (Extended Data Fig. 7 and Methods).

**Fig. 6.**
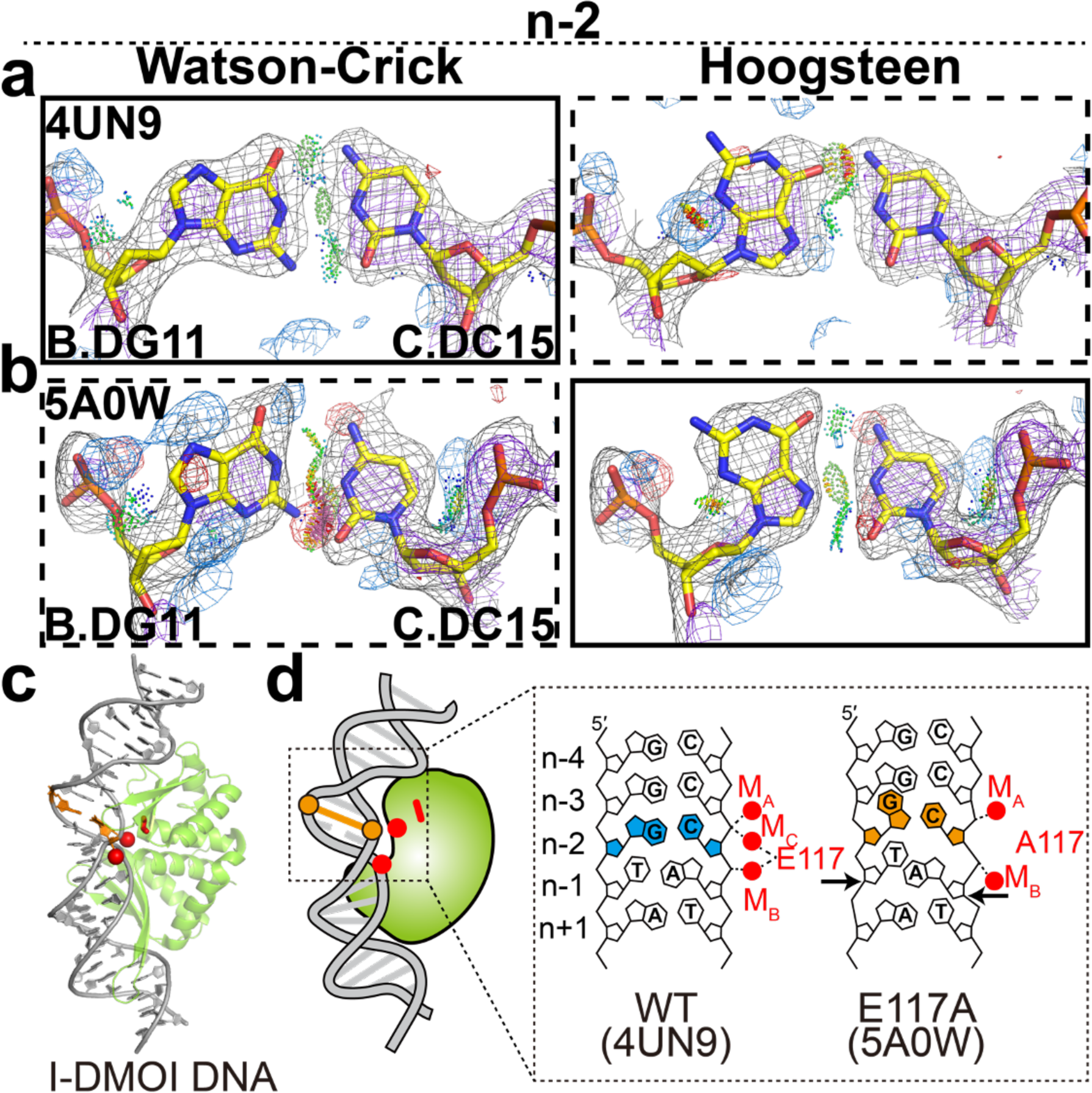
Hoogsteen base pairs in I-DMOI endonuclease. **(a,b)** Comparison of 2mF_o_-DF_c_ and mF_o_-DF_c_ electron density maps for the original Watson-Crick (left) and the corresponding Hoogsteen models (right) for the G-C bp at position *n*-2 in **(a)** a wild-type substrate complex, **(b)** the E117A mutant substrate complex. Electron density meshes and stereochemistry are as described in Fig. 1c and the box scheme is as described in Fig. 2a. **(c)** 3D structures of the protein-DNA complex showing the Hoogsteen bps. **(d)** Schematic showing the DNA containing Hoogsteen bps (in orange), Watson-Crick bps (in skyblue), as well as the E117/A117 (in red). Also shown are the metal ions (red filled circle).

**Fig. 7.**
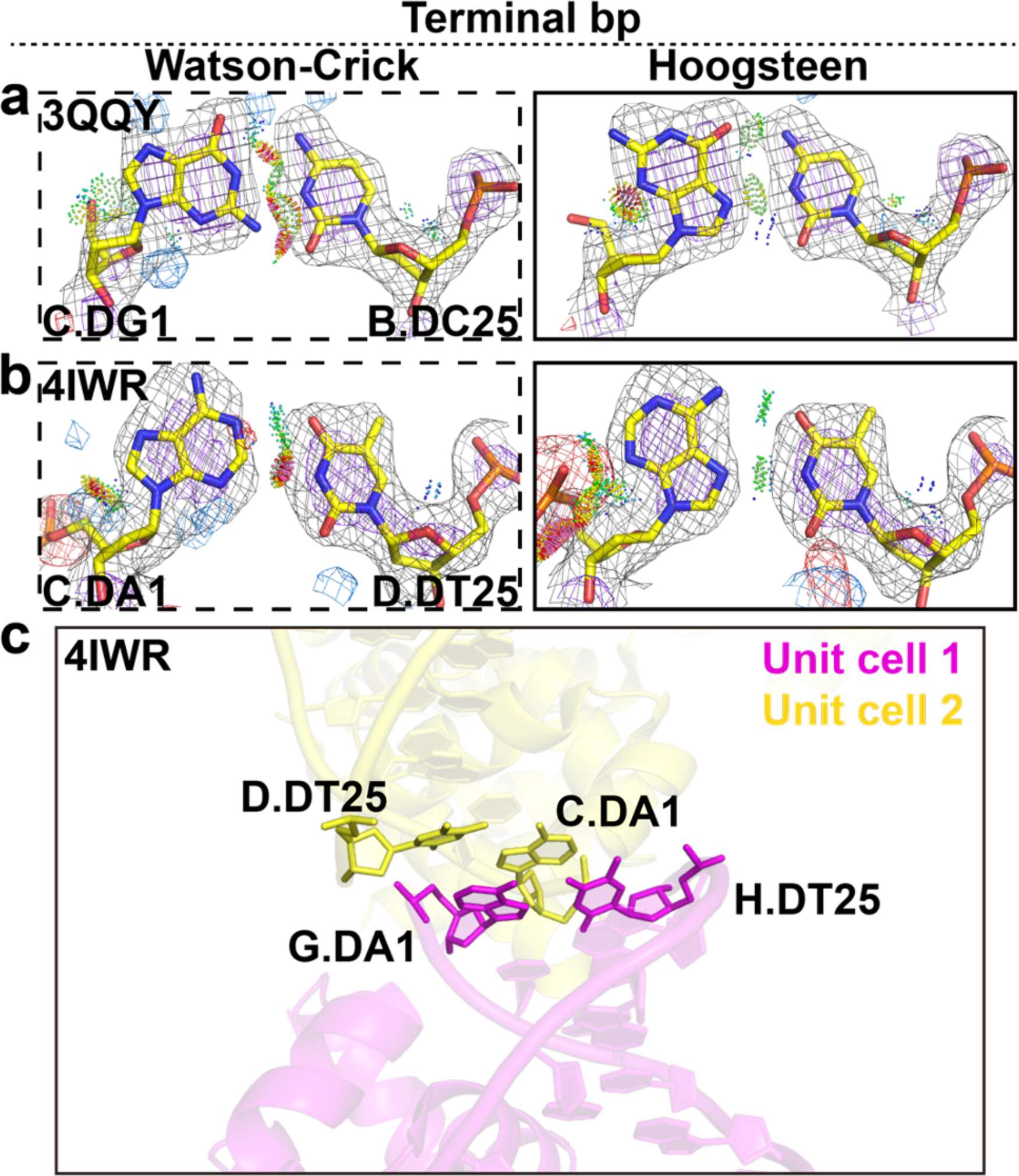
Terminal Hoogsteen base pairs. **(a,b)** Comparison of 2mF_o_-DF_c_ and mF_o_-DF_c_ electron density maps for the Watson-Crick (left) and the corresponding Hoogsteen models (right) for **(a)** a G-C terminal bp in Homing endonuclease I-Onul and **(b)** a A-T terminal bp in Esp1396I. Electron density meshes and stereochemistry are as described in Fig. 1c and the box scheme is as described in Fig. 2a. A complete set of data is provided in Extended Data Fig. 6-7. **(c)** An example of crystal stacking interactions in a terminal Hoogsteen bp.

As a positive control, our pipeline correctly uncovered all Hoogsteen bps that were mismodeled as Watson-Crick within the same position in the four complexes in the crystallographic ASU that were also identified in a prior study^20^ (Supplementary Tables 5, 6). In addition, several bps annotated as Hoogsteen or ambiguous Hoogsteen were found in structures of proteins previously shown to bind DNA in a Hoogsteen conformation (Extended Data Fig. 6-7, Supplementary Tables 5-7 and Supplementary Note 3-4). These include the tumor suppressor p53^7^ and *Sulfolobus solfataricus* polymerase Dpo4^10^.

In summary, ∼10% (n=22) of the bps identified using our pipeline were Hoogsteen, ∼63% (n=135) were ambiguous (including those with weak density), and only ∼27% (n=58) were Watson-Crick. The percentages of Hoogsteen (n=17, ∼9%), Watson-Crick (n=52, ∼26%), and ambiguous (n=130, ∼65% with n=21 ambiguous Hoogsteen) bps did not change substantially when curating the data to account for redundant bps (Fig. 1f, Supplementary Table 3 and Methods). As expected, we didn’t observe substantial differences between the R-work/R-free (R-factors) in the structures when they were refined with a Watson-Crick or Hoogsteen conformation (Supplementary Table 5). This underscores the limitations of R-factors in differentiating model differences that comprise a small percentage of the total structure. These results suggest there may be widespread ambiguities regarding the nature of base pairing in existing structures of DNA that are not well-documented, and that Hoog-finder provides a means for effectively identifying such bps.

### Hoogsteen bps are located near stressed DNA sites

Most of the newly identified Hoogsteen bps were located in stressed regions of DNA duplexes, which we define to be bps that are not flanked by canonical Watson-Crick bps as detected using X3DNA-DSSR^31^. Among the 17 Hoogsteen bps, 13 A(*syn*)-T and four G(*syn*)-C^+^, 16 (94%) were at or near regions of chemical and/or structural stress. Two were found next to a mismatch, two near lesions, two next to a nick, one near a melted bp, and nine were terminal bps (Fig. 1f and Supplementary Table 6). For comparison, only ∼20% of the total bps in the *Parent* dataset (∼100,000 bps) were in stressed regions of DNA duplexes.

### Hoogsteen base pairs near lesions

Among the eight non-terminal Hoogsteen bps, three were in crystal structures of DNA bound to the low fidelity polymerase *Sulfolobus solfataricus* polymerase Dpo4^32^ (Fig. 2a-c, Extended Data Fig. 6, Supplementary Table 6 and Supplementary Note 3). Fig. 2a shows the improvement in the electron density observed with the Hoogsteen versus Watson-Crick model in some cases accompanied by better stereochemistry including reduced steric clashes and more favorable H-bonding. An additional five ambiguous Hoogsteen bps in Dpo4 structures were also identified that showed a slightly better fit to the electron density when modeled as Hoogsteen relative to Watson-Crick and also showed some improvement in stereochemistry (Extended Data Fig. 7, Supplementary Table 7 and Supplementary Note 3). The large number of crystal structures available for Dpo4-DNA complexes (n=162) provided a unique opportunity to assess the role of DNA stress, in this case lesions and mismatches, in determining preferences for a Hoogsteen versus the Watson-Crick conformation. To aid this statistical analysis, we also considered those five ambiguous bps in Dpo4-DNA structures that show a slight preference for the Hoogsteen conformation.

Prior studies have identified Hoogsteen bps in crystal structures of Dpo4 in which they were proposed to accommodate lesion induced DNA distortions^10, 11, 22, 23^ to allow bypass of damage during replication^33^. Our new findings expand this Hoogsteen landscape, revealing Hoogsteen bps adjacent to a wider variety of damaged nucleotides (such as 2,4-difluorotoluene and S-methanocarba-dATP), sampling a broader variety of positions (*n*-3 in addition to the previously documented *n*-1 and *n*-2) relative to the active site, with two or as many as three consecutive Hoogsteen or ambiguous bps forming adjacent to one another (Fig. 2b-c and Supplementary Tables 6, 7).

Importantly, Hoogsteen bps were only observed in Dpo4-DNA crystal structures (n=9) with duplexes containing lesions or mismatches (Fig. 2c, Extended Data Fig. 9 and Supplementary Note 3). By contrast, 26 Dpo4-DNA crystal structures lacking lesions or mismatches were purely Watson-Crick (Supplementary Table 8). Not all structures, however, containing mismatches or lesions feature Hoogsteen or ambiguous bps. Instead, duplexes containing the lesions can be Hoogsteen, Watson-Crick or ambiguous bps depending on the identity of the base partner and/or position of lesion along the duplex (Fig. 2c and Extended Data Fig. 9).

As noted, Hoogsteen bps in the Dpo4-DNA structures tend to be observed at positions *n*-1 and *n*-2 near the active site (*n*) (Fig. 2c, Extended Data Fig. 9). Interestingly, we noticed that the C1′-C1′ distances at *n*-2 were slightly pre-constricted even when the bps are Watson-Crick in Dpo4 DNA lacking lesions or mismatches (Fig. 2d). Without the constriction at this position, steric collisions would occur with the Dpo4 protein (Fig. 2e). Thus, it appears that Dpo4 actively constricts the bp at this position, and that this in turn increases the propensity to form a Hoogsteen bp. A similar mechanism has been proposed to explain the preference of polymerase ι for Hoogsteen bps in its active site (*n*)^28^. These findings reinforce a prominent role for Hoogsteen bps in DNA damage and mismatch bypass by Dpo4. The Hoogsteen bps might serve to better absorb the conformational stress and deviation from canonical Watson-Crick geometry imposed by damaged nucleotides or mismatches.

### Hoogsteen base pairs near mismatches

We recently reported the first series of crystal structures for a transcription factor bound to a DNA duplex containing mismatches^30^. Although not discussed in the original publication, one of the structures (PDB: 6UEO) included a G(*syn*)-C^+^ Hoogsteen bps immediately adjacent to a partially melted A-C mismatch within the consensus sequence of TBP, a transcription factor shown previously to bind matched DNA in a Hoogsteen conformation^5^. The G(*syn*)-C^+^ Hoogsteen bp occurs at an unstacked step, an environment similar to duplex terminal ends, in which Hoogsteen bps are frequently found^16^ (Fig. 3a-c).

Interestingly, our new analysis identified other Hoogsteen bps next to mismatches, including two A(*syn*)-T Hoogsteen bps sandwiched between two C-T mismatches in a complex involving the endonuclease T5 flap (T5Fen). Here, the electron density and stereochemistry strongly favor the Hoogsteen over Watson-Crick model (Fig. 3d). This enzyme trims branched DNAs that arise from Okazaki-fragment synthesis^34^. The C-T mismatches were used to aid crystallization (PDB: 5HP4) in a region distant from the active site^35^ (Fig. 3d-f). Like Hoogsteen bps, pairing to form a C-T mismatch requires constriction of the two bases by ∼2.0-2.5 Å. Indeed, pre-constricted pyrimidine-pyrimidine mismatches such as C-T and T-T have recently been shown to mimic the distortions induced by Hoogsteen bps^30^. The T5Fen crystal structure suggests that in addition to structurally mimicking the constricted Hoogsteen conformation^30^, these mismatches can also promote Hoogsteen bps at neighboring sites.

We tested the above hypothesis for naked duplex DNA under solution conditions with the use of NMR relaxation dispersion (RD) experiments^36–38^. Strikingly, the equilibrium A-T Hoogsteen population increased by 3-fold and 13-fold when placed next to G-T and T-T mismatches, respectively (Fig. 3g-h and Extended Data Fig. 10). The larger boost in the Hoogsteen population seen adjacent to the constricted T-T mismatch relative to the unconstricted G-T wobble is consistent with constriction being an important force increasing the preference for Hoogsteen, though we cannot rule out other causes such as stacking.

### Hoogsteen base pairs near nicks

Nicked DNA is a form of damage and reaction intermediate that various enzymes act upon during DNA replication, damage repair, and gene editing^39–41^. Our pipeline identified Hoogsteen bps near nicked sites in crystal structures of DNA duplexes bound to two different proteins, human AP endonuclease 1 (APE1) and *Thermus thermophilus* Argonaute (TtAgo).

APE1 is a multifunctional enzyme. One of its roles is an exonuclease removing 3′ lesions^42, 43^ to enable downstream repair. Through its exonuclease activity, APE1 is proposed to help proofread polymerase β insertions during BER by removing mis-inserted bases to regenerate a gapped DNA^44–46^. In this role, APE1 needs to act on the mis-inserted mismatched base adjacent to a 3′ nick while discriminating against a correctly inserted Watson-Crick bp.

In the crystal structure (PDB: 5WN4) of the catalytically active substrate complex of APE1 bound to a nicked DNA duplex containing template thymine and mis-inserted cytosine, the T-C mismatch within the active site is melted and the DNA backbone is sharply bent within the catalytic pocket (Fig. 4a, d-e). The *n*-1 and *n*-2 bps adjacent to the mismatch have weak electron density and we annotate them as ambiguous (Fig. 4a). A similar structure was observed (PDB: 5WN1) for the product complex following excision of the mis-inserted cytosine in which positions *n*-1 and *n*-2 form well-resolved Watson-Crick bps (Fig. 4b, d-e).

In contrast, in the corresponding APE1-DNA crystal structure (PDB: 5WN0) with template guanine, the correctly inserted cytosine formed the expected Watson-Crick G-C bp (Fig. 4d-e). However, the structure of this complex differs substantially from that of the T-C mismatch. The inserted cytosine is displaced 7.5 Å away from the active site. In the original publication^46^, this inactive APE1-DNA conformation was proposed to explain how it discriminates and avoids cleaving matched Watson-Crick DNA.

Interestingly, our analysis identifies the *n*-1 and *n*-2 bps in this inactive structure to be A(*syn*)-T and G(*syn*)-C^+^ Hoogsteen bps, respectively (Fig. 4c). The electron density here is not as strong as for some of the other examples and it is clearer for the position *n*-1 versus *n*-2, with both positions showing improved stereochemistry with the Hoogsteen model (Fig. 4c). By unwinding the DNA ∼12°, the Hoogsteen bps appear to induce a register shift so that they now occupy the active site in place of the G-C Watson-Crick bp, displacing the inserted cytosine away from the active site (Fig. 4d-e and Supplementary Table 9). In addition, one of the key catalytic residues, Arg177 is recruited to the Hoogsteen bps where it stacks on the thymine base and forms H-bonds with the thymine phosphate backbone at position *n*-1. Notably, position *n*-1 was also shown to be a Hoogsteen bp in Fig. 4b of the original publication by Whitaker *et al*^46^; however the bp is modeled as Watson-Crick in the deposited PDB and no reference was made to the Hoogsteen bp in the publication^46^. The Hoogsteen bps may help increase the specificity of APE1 through an induced-fit^47^ mechanism by stabilizing a catalytically inactive conformation when bound to a matched Watson-Crick bp.

Our analysis also identified an A(*syn*)-T Hoogsteen bp near a nick in crystal structures (PDB: 4KPY, 4NCA, 4NCB) of the TtAgo-DNA complex. TtAgo employs short 13-25 nt single-stranded DNA guides to introduce nicks between positions *n* and *n*+1 in single-stranded RNA during RNA silencing^48, 49^ and in single-stranded DNA as part of a defense system^49, 50^. Prior crystal structures of TtAgo complexes with guide and target DNA revealed a transition between inactive and active conformations that ensures specificity toward substrates of specific length. During this transition, the highly conserved catalytic residue Glu512 moves near the binding pocket where it contacts the DNA backbone at position *n*+4, forming water mediated contacts with catalytic metal ions^51^.

In both the inactive (PDB: 4N41) and the active (PDB: 5GQ9) substrate complex (PDB: 5GQ9) where the target DNA is not cleaved, the bp at position *n*+4 is more favored as Watson-Crick bp (Fig. 5a-b, d-e). However, in a crystal structure (PDB: 4KPY) of a product complex where the target DNA is cleaved (with DNA nicked between *n* and *n*+1), our analysis indicates that the bp at position *n*+4 is an A(*syn*)-T Hoogsteen bp (Fig. 5c). Here, the strong positive difference densities around adenine N7/C5/N6 and at N3 in the Watson-Crick conformation essentially disappear when the base is modeled and refined in the Hoogsteen conformation (Fig. 5c). The Hoogsteen bp retains the same contacts with Glu512 as observed in the Watson-Crick conformation (Fig. 5e). Although it remains unclear what interactions favor the Hoogsteen bp, the density at this position also slightly favors the Hoogsteen conformation in two other related crystal structures (PDB: 4NCA and 4NCB) (Extended Data Fig. 7 and Supplementary Table 7). Moreover, a preference to form a Hoogsteen bp at position *n*+4 was robustly observed for the same bps in complexes that were present as multiple copies in the crystallographic asymmetric unit (ASU). Therefore, these data are suggestive of a Watson-Crick to Hoogsteen transition taking place during the catalytic cycle, but this requires further investigation.

Our analysis also identified an ambiguous A(*syn*)-T Hoogsteen bp adjacent to a nick in the crystal structure of an inactive hairpin-forming complex of the RAG1/2 recombinase (PDB: 5ZDZ) (Extended Data Fig. 11 and Supplementary Table 7). Together with the prior crystal structure of the IHF-DNA complex^4^, these results suggest a preference for Hoogsteen bps adjacent to nicked sites.

### Hoogsteen bps involving interactions with metal ions

Our analysis also identified a Hoogsteen bp in a crystal structure of the homing endonuclease I-DMOI that appears to be stabilized through interactions with metal ions. I-DMOI sequence specifically recognizes and cleaves a stretch of 22 bps of double-stranded DNA^52^. In the crystal structure (PDB: 4UN9) of the catalytically active conformation, the DNA within the active site is locally overwound and has a substantially narrowed minor groove (Supplementary Table 9). The catalytic residue Glu117 forms contacts with two metals, termed M_B_ and M_C_, which in turn form a network of interactions with the DNA, stabilizing a strained conformation at positions *n*-1 to *n*-3 (Fig. 6a, c-d).

Interestingly, in the corresponding crystal structure (PDB: 5A0W) of a mutant of I-DMOI with Glu117 replaced by Ala117, M_C_ is no longer observed, while M_B_ changes coordination likely to compensate for loss of contacts with Glu117 (Fig. 6c-d). This change in metal coordination is accompanied by a change in the DNA conformational strain, particularly at position *n*-1. Rather than base pairing, the adenine and partner thymine stack on top of each other, thus constricting the DNA (Fig. 6d). Immediately adjacent to this unusual A/T stack at position *n*-2, our analysis identified a G(*syn*)-C^+^ Hoogsteen bp, which may help to absorb the unusual constriction at the neighboring position *n*-1 (Fig. 6b). Here, the electron density and stereochemistry very clearly favor the Hoogsteen over the Watson-Crick model (Fig. 6b). As proposed in the original paper^53^, it is possible that the newly positioned metal M_B_, and phosphate group stabilizes this new type of strain. The same bp in the other complexes in the ASU were also identified as Hoogsteen.

It is noteworthy that the ambiguous Hoogsteen bp observed next to a nick in the crystal structure of the inactive hairpin-forming complex of the RAG1/2 recombinase (PDB: 5ZDZ) also featured changes in metal coordination to the DNA relative to the active Watson-Crick form (Extended Data Fig. 11, Supplementary Table 7 and Supplementary Note 5), providing an additional example in which metals appear to participate in Hoogsteen bp formation.

### Terminal Hoogsteen bps

Many biochemical processes act on the terminal ends of DNA duplexes, including homologous recombination and nonhomologous end joining^54^. There is evidence showing a preference for Hoogsteen bps to form within terminal ends of DNA duplexes. The prior Hoogsteen survey^16^ identified at least 10 A(*syn*)-T and two G(*syn*)-C^+^ terminal Hoogsteen bps distant from the protein binding site in 10 crystal structures of protein-DNA complexes. In addition, Hintze *et al.*^20^ identified an additional 4 A(*syn*)-T and two G(*syn*)-C^+^ Hoogsteen bps that were mismodeled as Watson-Crick, also distant from the protein binding site. As noted by Hintze *et al.*^20^, some of these terminal Hoogsteen bps could be stabilized by crystal contacts. However, solution state NMR RD studies also show a 4-fold higher propensity to form Hoogsteen bps at DNA terminal ends relative to the center of a DNA duplex^55^.

Our current analysis uncovered two new terminal Hoogsteen bps positioned also distant from protein binding sites (Supplementary Table 6). These include a G(*syn*)-C^+^ bp in the DNA of a homing endonuclease I-Onul complex (PDB: 3QQY) (Fig. 7a), and an A(*syn*)-T bp in the DNA complexed with the regulatory protein Esp1396I of the type II restriction-modification (RM) system (PDB: 4IWR) (Fig. 7b). Here, both the electron density and stereochemistry favor the Hoogsteen over the Watson-Crick model (Fig. 7a-b). We also identified several ambiguous Hoogsteen bps at DNA terminal ends (Extended Data Fig. 7, Supplementary Table 7 and Supplementary Note 6). We cannot rule out that these terminal Hoogsteen bps are induced by crystal contacts, as all of them are involved in packing with neighboring symmetry related molecules in the crystal unit cell. For example, the terminal Hoogsteen bp in Esp1396I stacks with a symmetry related Hoogsteen bp from a neighboring complex in the crystal (Fig. 7c).

## Discussion

It is commonly assumed that A-T and G-C bps in duplex DNA are Watson-Crick. However, prior studies showed that certain proteins^4–7^ and drugs^12^ bind to specific DNA sequences and render the Hoogsteen bp as the dominant conformation at certain positions. Our results suggest that Hoogsteen bps are not restricted to a few transcription factors or specialized polymerases, but may in fact be a more common feature of conformationally stressed DNA, also found in complexes with enzymes that repair or cleave DNA.

In particular, forms of stress that result in the constriction of the helical diameter, such as pyrimidine-pyrimidine mismatches and stacking of base partners, or that result in an environments mimicking the terminal ends, such as nicks, appear to favor the Hoogsteen conformation. Interestingly, in the crystal structure of the IHF-DNA complex^4^, a Hoogsteen bp was only observed at the nicked site but a Watson-Crick bp was observed at a symmetrically pseudo-symmetry related site lacking the nick. In addition, solution NMR studies^56^ revealed that the Hoogsteen bp observed in the crystal structure of the complex does not form in an intact DNA duplex lacking the nick. Thus, the Hoogsteen bp observed in the IHF-DNA complex can directly be attributed to the nick.

The crystallographic and NMR evidence presented here showing a preference for Hoogsteen bps near mismatches is of particular interest considering a recent study^30^ showing that introducing mismatches, including pyrimidine-pyrimidine mismatches that favor Hoogsteen bps at specific positions in duplex DNA, can increase transcription factor binding affinity. High affinity transcription factor binding to mismatched DNA could compete with damage repair and promote mutagenesis at transcription factor binding sites^57^. The increased binding affinity imparted by mismatches was previously attributed in part to pre-paying the energetic cost of deforming the DNA for protein recognition. Based on our results, Hoogsteen bps near mismatches could also contribute to high affinity binding to mismatched DNA.

Even for the bps annotated as Hoogsteen, the weight of the crystallographic evidence varied from case to case. Whether these newly uncovered Hoogsteen bps also form under physiological solution conditions remains to be established. It will therefore be important to apply complementary solution-state approaches to test the validity of these Hoogsteen bps, resolve ambiguous bps, and also provide insights into any Watson-Crick to Hoogsteen dynamics that may be taking place. In the case of p53-DNA complex, the tandem A(*syn*)-T Hoogsteen bps observed in crystal structures^7^ could be verified independently under solution conditions using chemical substitutions^29^ and more recently, via high throughput binding measurements^30^. Similarly, the G(*syn*)-C^+^ Hoogsteen bps observed in crystal structures of TBP^5^ were recently verified under solution conditions using IR spectroscopy^58^. These and other chemical probing approaches^59^ could be used to verify the newly indentidied Hoogseen bps under solution conditions.

While we have proposed potential roles for some of the newly identified Hoogsteen bps, future studies could more directly examine their biological significance. Here, approaches similar to those first introduced to study the iota polymerase^9, 60^ could be applied: one examines how deazapurine substitutions which selectively destabilize the Hoogsteen bp^61^, or pyrimidine-pyrimidine substitutions which mimic the Hoogsteen bp^30^, impact binding affinity and/or enzymatic activity.

There is good reason to believe that additional Hoogsteen bps remain to be uncovered that are presently modeled as Watson-Crick in existing crystal structures of DNA. Our pipeline only analyzed the electron density for ∼200 out of ∼90,000 bps satisfying all three positive structural fingerprints, yet based on our training dataset, we know that some Hoogsteen bps only satisfy a subset of the criteria. There are an additional ∼1,400 bps that remain to be analyzed that satisfy the key C1′-C1′ distance criteria, which appears to be the most reliable diagnostic feature of a Hoogsteen conformation. In addition, Hoog-finder will likely fail to identify Hoogsteen-like conformations found in a previous survey of crystal structures^16^, in which the two base partners are not constricted but form H-bonds with *syn* purine bases.

Equally importantly, many of the DNA bps analyzed in existing crystal structures could not be definitively modeled as either Watson-Crick or Hoogsteen. Among the ∼200 bps satisfying all three structural fingerprints, over 60% were ambiguous. In this regard it is notable that our data indicate that Hoogsteen bps tend to be located adjacent to mismatches, lesions or nicks, which may be more flexible. However, it remains to be seen whether the weakening of electron density at some of these sites originates from increased flexibility. Future studies should also explore the application of ensemble-based refinement of both Watson-Crick and Hoogsteen models with fractional populations^62, 63^. Together with prior studies showing the ambiguity when modeling Hoogsteen versus Watson-Crick^7, 8, 18–20^, these results underscore the importance of exercising caution when modeling DNA bases, test Hoogsteen and other conformational states as a possible alternative, and annotate those bps that have ambiguous electron density.

Our approach identified 13 new Hoogsteen bps (Supplementary Table 6), which were not previously identified in the study by Hintze *et al.*^20^, which utilized as the sole diagnostic, the pattern of difference electron density peaks (Fig. 1b). Indeed, some of the bps, which we found to be mismodeled as Watson-Crick but are really Hoogsteen, did not show all the expected diagnostic difference electron density peaks used by Hintze *et al.*^20^ (Supplementary Note 1). However, this is not surprising given the relatively low resolutions of some of the structures and/or the weak electron density in the vicinity of the given bps; hence the difference densities in some of the structures were not highly reliable. In fact, half of Hoogsteen bps mismodeled as Watson-Crick in the *training* dataset lack the precise diagnostic difference density peaks and therefore could not be identified by the *find_purine_decoy* program developed in Hintze *et al.*^20^ (Extended Data Fig. 1 and Supplementary Note 1). Future studies could combine aspects of the two approaches to most effectively flag for potentially Hoogsteen bps mismodeled as Watson-Crick.

Finally, we hope that these findings will help spur a community-wide effort to re-analyze existing structures of DNA to consider the possibility of Hoogsteen and perhaps other bp conformations and to find ways to resolve bp conformation ambiguities in crystal structures and to also consider the Hoogsteen conformation when solving future crystal structures of DNA.

## Acknowledgments

We thank members of the Al-Hashimi laboratory for assistance and critical comments on the manuscript, Prof. Zippora Shakked (Weizmann Institute of Science) for bringing to our attention the Hoogsteen base pair in the crystal structure of T5 flap endonuclease (PDB: 5HP4), Prof. Mark Wilson (University of Nebraska) for critical input during early stages of the project, and Dr. Bradley Hintze (Duke University) for assistance.

## Author contributions

H.S., M.A.S. and H.M.A. conceived the project and experimental design. H.S. performed the structural survey. H.S. performed X-ray structure refinement and other structural analysis, with assistance from M.A.S., H.-F.L. and U.P. I.J.K. prepared NMR samples, performed NMR experiments and analyzed NMR data. H.M.A. and H.S. wrote the manuscript with critical input from M.A.S.

## Methods

### Generating *training* dataset of Hoogsteen bps mismodeled as Watson-Crick

The *training* dataset (n=28) was generated based on a previous X-ray structural survey of Hoogsteen bps^16^. We selected all the non-redundant Hoogsteen bps from Table 1 in Zhou *et al.*^16^, excluding structures with no deposited structure factors (e.g. Triostin A-DNA complex, PDB: 1VS2), with multiple models (e.g. terminal bps in Echinomycin-DNA complex, PDB: 1XVN), or with modified purine bases (e.g. the m^1^A(*syn*)-T bp in ALKBH2-DNA complex, PDB: 3H8O). To this dataset we also added two recent examples of G(*syn*)-C^+^ Hoogsteen bps from two recently solved crystal structures of the TBP-DNA complex (PDB: 6NJQ, 6UEO), which were not included in Zhou *et al.*^16^. The final dataset contained a total of 28 Hoogsteen bps (22 A(*syn*)-T and 6 G(*syn*)-C^+^ Hoogsteen) (Supplementary Table 1).

All the Hoogsteen bps in the *training* dataset were then mismodeled as Watson-Crick bps using the following procedure: (1) The coordinates for the *syn* purine residue in the Hoogsteen bp was removed from the original coordinate file. (2) An *omit* map of the original coordinate file was then generated by refining the coordinates in *phenix.refine*^25^ using the default settings in the *PHENIX* software^24^. (3) An *anti* purine residue was modeled into the resulting *omit* map and optimized via real space refinement using *COOT*^64^. (4) A second round of refinement was conducted using the same *phenix.refine* routine with the remodeled coordinates. The stereochemistry of different bp models were assessed using *MolProbity*^26^.

### Identification of structural fingerprints for the *training* dataset

X3DNA-DSSR^31^ was used to analyze all the nucleotide torsion angles (α, β, γ, δ, ε, ζ, χ, sugar phase angle) as well as all the bp parameters (shear, stretch, stagger, buckle, propeller twist, opening, C1′-C1′ distance) of the mismodeled Watson-Crick bps in the *training* dataset (n=28) as well as for canonical Watson-Crick bps (n=149) from a previous structural survey^16^. The sign of the raw output values for bp parameters shear and buckle were adjusted according to the index order of purine and pyrimidine as described in^30^.

### Generating the *negative training* dataset of Watson-Crick bps mismodeled as Hoogsteen

The negative *training* dataset (n=10) was generated by selecting a subset of the well-resolved Watson-Crick bps (five A-T and five G-C bps) from the canonical Watson-Crick bps (n=149) from Zhou *et al.*^16^ (Supplementary Table 2). We performed a similar procedure as described in **Generation of a *training* dataset**, but this time we flipped the *anti* purine to be the *syn* conformation and followed the same refinement protocol. The bp parameters (shear, stretch, stagger, buckle, propeller twist, opening) cannot be interpreted because they are ill-defined for the Hoogsteen bp given a change in the coordinate reference frame as described previously^30^.

### Screening putative Hoogsteen candidates in X-ray structures using Hoog-finder

X-ray structures of protein-DNA complexes (defined as PDB structures with both DNA and protein present in the macromolecular entities) with resolution ≤ 3.5 Å were downloaded as cif files with available electron density map coefficients from the RCSB website (www.rcsb.org) on Aug 29^th^ 2020. For palindromic DNA that were deposited as single chains in the ASU, the biological assemblies containing the double stranded models were downloaded from RCSB and processed by X3DNA-DSSR with the symmetry flag “--symm”. X3DNA-DSSR was then used to parse the structural descriptors of bps from all PDB structures into a searchable database, which included nucleotide local torsion angles (α, β, γ, δ, ε, ζ, χ, sugar phase angle), bp parameters (shear, stretch, stagger, buckle, propeller, opening), C1′-C1′ distance. We then searched for potential candidate bps that were Hoogsteen but mismodeled as Watson-Crick based on the following the queries:

1. We only considered dA-dT or dG-dC bp with Watson-Crick geometry defined by the *Leontis-Westhof* (LW) notation as “cWW”, “cWS”, “cW.”, which excluded all the *trans* bps, Hoogsteen bps or any platform bps.
2. Based on the structural fingerprints of mismodeled Watson-Crick bps in the *training* dataset, we only considered bps that satisfy shear > 0.5 Å, opening > 10° and C1′-C1′ distance < 10.0 Å simultaneously.
3. We manually checked and excluded cases including bps from tertiary interactions, misaligned bps that are false positive, bps in DNA regions with potential two-fold statistical disorder, bps with multiple modeling, and identical bps due to crystal symmetry.

Among 97,100 A-T and G-C Watson-Crick bps (*Parent* dataset, n=97,100), there are a total of 215 bps satisfying all the queries described above (*Starting dataset*, n=215) (Fig. 1e and Supplementary Tables 3, 4). We then manually inspected the local electron density of the bps and further removed cases of either weak local density which are difficult to model any bp (n=91) or well-resolved Watson-Crick density (n=58) to yield a smaller dataset (*Filtered* dataset, n=66) (Fig. 1e, Extended Data Fig. 4a-b, Supplementary Tables 3-4). These 66 candidates were then subjected to a similar procedure in **Generation of a *training* dataset**, but this time we flipped the *anti* purine to the *syn* conformation to generate Hoogsteen bps for structural refinement. We then compared the agreement of the electron density as well as the improvement of stereochemistry of the two bases between the Watson-Crick and the Hoogsteen model. As a result, 22 bps were identified as Hoogsteen bps which showed better improved agreement with the electron density and the stereochemistry and didn’t resemble the distorted Hoogsteen geometry in the *negative training* dataset. The remaining 44 bps were denoted as ambiguous bps as we were not able to confidently discriminate Watson-Crick versus Hoogsteen (Extended Data Fig. 4c and Supplementary Note 2). Among those 44 ambiguous bps, 23 bps are slightly favored as Hoogsteen than Watson-Crick (‘ambiguous Hoogsteen’) (Extended Data Fig. 7). Compared to the Hoogsteen bps, the improvements in modeling the electron density and/or stereochemistry for Hoogsteen relative to Watson-Crick bps were not as substantial for these 23 ambiguous Hoogsteen bps. In total, there were 22 Hoogsteen (10%) and 58 Watson-Crick (27%) in the *Starting* dataset, and the remaining 135 were ambiguous bps (63%) including those with weak local electron density.

In the structures where we found Hoogsteen or ambiguous Hoogsteen bps, there are sometimes more than one repeating protein-DNA complex within a single ASU. However, not all the bp positions were identified by our structure-based screening. This is either because they did not form H-bonds detectable by 3DNA which were not included in the *Parent* dataset or because they failed to satisfy all three criteria applied due to subtle structural differences between different protein-DNA complexes (Supplementary Table 4). Therefore, we manually analyzed the electron densities of additional bps which were not identified by the structure-based screening. Indeed, we found four more Hoogsteen bps and seven more ambiguous Hoogsteen bps (Supplementary Table 4).

The bps at the same positions as those in the other protein-DNA complexes in the ASU were considered as redundant bps and were subsequently removed from the curated dataset. The final result after these data curation can be summarized as 17 Hoogsteen (9%), 52 Watson-Crick (26%) and 130 ambiguous (65%) bps with 21 ambiguous Hoogsteen bps (Fig. 1f and Supplementary Table 3).

A similar analysis was also carried out for structures of DNA without protein bound but no putative Hoogsteen bps emerged.

### Structural analysis

#### DNA global shape

DNA major and minor groove widths were quantified by the P-P distance metric^65^ using X3DNA-DSSR^31^. DNA inter-helical local kinking and twisting were quantified by a Euler angle approach as described before^16^. In this approach, two 2-bp idealized B-form DNA helices (H1 and H2) were generated by 3DNA^66^ and were superimposed on the DNA structure immediately above and below a specific junction (J) bp. The H1 is specified by the 5′-direction of one of the J residues (in “nt_1” columns in Supplementary Table 9). The resulting orientation of the H1 and H2 was then calculated using three inter-helical Euler angles (α_h_, β_h_, γ_h_) relative to a reference helix, in which the two helices are coaxially aligned in an idealized B-form helix geometry^16^. The inter-helical Euler angle β_h_ (0°≤β_h_≤180°) therefore defines the local kink angle about the J bp, while γ_h_ (−180°≤γ_h_≤180°) defines the directionality of kinking, with γ_h_ = +/-90° indicating major groove and γ_h_ = −180°≤γ_h_≤-90° or 90°≤γ_h_≤180° indicating minor groove directed kinking, respectively^16^. The inter-helical twist angle ζ_h_ = α_h_ + γ_h_ describes the relative twist between H1 and H2 with ζ_h_ > 0° and ζ_h_ < 0° representing over- and unwinding, respectively^16^. All the calculations with poor alignment to the idealized B-form helix (RMSD > 2 Å using all backbone atoms) were excluded as poor agreement to an idealized helix leads to unreliable Euler angles.

#### DNA protein interactions

H-bonding and van der Waals interactions between DNA and protein were detected by a web-based tool: DNAproDB^67^ (https://dnaprodb.usc.edu/index.html).

### NMR experiments

All the DNA constructs (hpCG, hpTA, hpTG, hpTT) used for NMR *R*_1ρ_ measurements are summarized in Extended Data Fig. 10a. ^13^C,^15^N uniformly labeled DNA samples were synthesized following the procedure described in Zimmer and Crothers, 1995^68^. The buffer used for NMR measurement was composed of 25 mM NaCl, 15 mM Na_3_PO_4_, 0.1 mM EDTA, 10% D_2_O at pH 5.9.

NMR ^13^C *R*_1ρ_ experiments were carried out on Bruker Avance III 600 MHz equipped with a triple-resonance HCN cryogenic probe as described previously^69^. Resonance assignments for hpTG were reported previously^69^ while assignments for other constructs were readily obtained by overlaying spectra to the hpTG construct. The spinlock powers and offsets used in the *R*_1ρ_ experiments were summarized in Supplementary Table 10. The analysis of *R*_1ρ_ data was also described in a prior study^69^. The fitting parameters of all the *R*_1ρ_ profiles were listed in Supplementary Table 11.

## Data Availability

The PDB coordinate files and structure factor amplitudes (MTZ) files of all the structural models rebuilt and refined with Hoogsteen bps in our study can be downloaded from: https://github.com/alhashimilab/HoogsteenInTheData. All other data supporting the findings are available within the article and its supplementary information.

