## Supplementary Information for "Revealing A-T and G-C Hoogsteen base pairs in stressed protein-bound duplex DNA"

**Supplementary Note**

**Supplementary Note 1: Electron density patterns of Hoogsteen bps mismodeled as Watson-Crick in *training* dataset**

In a previous study^1^, Hoogsteen bps mismodeled as Watson-Crick were identified by screening coordinate/electron density maps for the presence of negative electron density at the purine N1/C2/C8 atoms and positive electron density at the purine N7/C4 atoms. In total, 15 out of 28 bps in our *training* dataset resulted in the expected diagnostic electron density patterns as described by Hintze *et al.* (Extended Data Fig. 1). Interestingly, the remaining 13 bps reveal different electron density patterns and therefore couldn't be identified by the *find_purine_decoy* program developed by Hintze *et al.* (Extended Data Fig. 1). For example, mismodeling the G(*syn*)-C^+^ Hoogsteen bp in the TBP-DNA structure (PDB: 6NJQ) as a Watson-Crick bp resulted in positive density near the purine N3 and negative density near the purine O6. In the case of the terminal Hoogsteen bps in the prototypical Apicomplexan Apetala2 DNA complex (PDB 3IGM), mismodeling the A(*syn*)-T Hoogsteen bp as a Watson-Crick bp resulted in minimal difference density peaks but caused a severe steric clash between the adenine and thymine as detected by *MolProbity*^2^.

Our study identified 13 additional Hoogsteen bps that were mismodeled as Watson-Crick relative to Hintze *et al.* (Supplementary Table 6). This is likely due to (1) inclusion of new PDB entries between June 2014 and Sep 2020 (2) the fact that some bps (e.g. the terminal bp in PDB: 3QQY in Fig. 7a) do not show the precise diagnostic patterns of difference electron density peaks used by Hintze *et al.* to identify potential Hoogsteen bps, similar to the *training* dataset mentioned above.

**Supplementary Note 2: Ambiguous examples of discriminating Watson-Crick and Hoogsteen**

Our study identified a total of 130 non-redundant bps that are ambiguous. The vast majority of these bps (n=91) exhibit weak local electron density which prevented clear modeling of the bp conformation (Extended Data Fig. 4b and Supplementary Table 4). For 14 of the remaining ambiguous bps (Supplementary Table 4), we are still unable to clearly differentiate between Watson-crick versus Hoogsteen based on electron density and stereochemistry. In some cases, both Watson-Crick and Hoogsteen bps showed nearly identical stereochemistry and agreement with the local electron density. For example, modeling the A-T bp at the 5′-end of the PAM sequence in the DNA in the CRISPR-Cas9 complex (PDB: 5X2G) as either Watson-Crick or Hoogsteen resulted in similar stereochemistry and agreement to the electron density (Extended Data Fig. 4c). As another example, modeling the A-T bp at position *n*-1 in the mouse polymerase β(I260Q)-DNA complex (PDB: 3UXP) as either Watson-Crick or Hoogsteen resulted in unfavorable steric clashes with similar agreement to the electron density (Extended Data Fig. 4c). In some cases, the local electron density is strong enough to model a bp but still too weak to discriminate between a Watson-Crick versus a Hoogsteen conformation. An example is the helical bp in the DNA bound to the Streptomyces transcriptional factor CprB (PDB: 4PXI) (Extended Data Fig. 4c). These examples illustrate the various types of ambiguous bps that were found in several protein-DNA structures.

**Supplementary Note 3: Hoogsteen bps in Polymerase Dpo4**

Polymerase Dpo4 is a low-fidelity polymerase that can bypass DNA lesions^3^. Hoogsteen bps have previously been observed in Dpo4 DNA complexes either at or adjacent to the polymerase active site. In one of these crystal structures (PDB: 1RYS), an A-T Hoogsteen bp was observed at the polymerase active site (position *n*) where the T forms a *cis-trans* thymine dimer with the adjacent T in the template strand. This A-T Hoogsteen bp was proposed to lubricate the DNA backbone distortions induced by the thymine dimer and provide a mechanism to bypass UV damage^4^ (Extended Data Fig. 9). In a second Dpo4-DNA crystal structure (PDB: 1S0M), an A-T Hoogsteen bp was also observed at the active site but next to a bulky Benzo[a]pyrene diol epoxide (BPDE)-adenine adduct paired with a T at position *n*-1 (Extended Data Fig. 9). Similarly, in a third Dpo4-DNA crystal structure (PDB: 2IBK), tandem A-T Hoogsteen bps were modeled at positions *n* and *n*-1 next to a DNA bulge with a bulky DNA damaged Benzo[a]pyrene-N^2^-dG (BP-dG) adduct between positions *n*-1 and *n*-2 (Extended Data Fig. 9). These Hoogsteen bps were previously proposed to accommodate the distortions due to the damaged bulge and allow the primer to fit into the active site^5,6^. In two more Dpo4-DNA crystal structures (PDB: 3V6J, 3V6H), there is a damaged guanine base with a bulky lesion N^2^,3-ethenoguanine (εG) in the insertion site (position *n*) and a G-C^+^ Hoogsteen bp was modeled at position *n*-1, which was proposed to enable better stacking with εG^7^ (Fig. 2c). Another G-C^+^ Hoogsteen bp was observed in crystal structure (PDB: 2W8K) containing a damaged G with a bulky adduct group Naphthyl at the Watson-Crick face, which destabilizes the Watson-Crick bp (Extended Data Fig. 9).

We identified three additional Hoogsteen bps in Dpo4-DNA crystal structures at DNA position *n*-2 (PDB: 1S97, 3V6J, 3V6H) and 5 additional ambiguous Hoogsteen bps at DNA positions *n*-1, *n*-2 and/or *n*-3 (PDB: 2VA2, 3V6J, 3V6H, 4GC7) (Fig. 2c and Supplementary Tables 6, 7), all of which are adjacent to the polymerase active site and surrounded by non-canonical bps. The bps near the active site (*n*-1, *n*-2) appear to be constricted by Dpo4 (Fig. 2d-e). In addition, in some structures, more extensive van der Waals contacts are observed between Dpo4 amino acid residues and the Hoogsteen *syn* purine base as compared to the corresponding Watson-Crick counterpart (Supplementary Table 12). These findings support a prominent role for Hoogsteen bps in DNA damage and mismatch bypass by Dpo4. The Hoogsteen bps might serve to better absorb the conformational stress and deviation from canonical Watson-Crick geometry imposed by damaged nucleotides or mismatches impinge.

**Supplementary Note 4: Ambiguous Hoogsteen bps in p53 and p73**

Prior studies have documented the propensities to form two adjacent (tandem) A(*syn*)-T Hoogsteen bps within the consensus sequence CWWG (W= A or T) of p53-DNA complexes^8^. Hoogsteen bps have been reported in 9 out of 29 crystal structures of p53-DNA complexes with available electron density. As was previously noted^9^, these tandem Hoogsteen bps with well-resolved electron density seem to be favored in cases where the consensus sequence is CATG in contiguous half-sites. In addition, a prior crystallographic analysis^8^ identified two tandem ambiguous Hoogsteen bps, which were originally modeled as Watson-Crick in a crystal structure (PDB: 3EXJ) of a mouse p53-DNA complex. In our study, we identified two additional tandem ambiguous Hoogsteen bps (Extended Data Fig. 7 and Supplementary Table 7) in the CATG bound to an artificially designed p53 stabilizing mutant (PDB: 3Q05), where all the protein mutations are far from the DNA. However, the corresponding tandem bps at the other half of the binding site which have the same conformation are better modeled as Watson-Crick. These findings could indicate a potential dynamic equilibrium between Watson-Crick and Hoogsteen bps in DNA sites bound by p53.

For the transcription factor p73, which has some structural similarity to p53, prior studies considered the possibility of having corresponding Hoogsteen bps at its DNA binding site^10^. Indeed, in the p73-DNA complex (PDB: 4G82), we identified one of the central tandem A-T bps as an ambiguous Hoogsteen bp (Extended Data Fig. 7 and Supplementary Table 7).

**Supplementary Note 5: Ambiguous Hoogsteen bps in recombinase RAG1/2**

The RAG1/2 recombinase initiates DNA recombination events by cleaving and splicing multiple immunoglobulin encoding DNA segments. This leads to the generation of immunoglobulin and T cell receptors that are critical for the vertebrate immune response^11^. To do this, a dimeric RAG1/2 sequence-specifically recognizes a pair of DNA sites, introduces a nick on one strand, and then simultaneously nicks the complementary strand at the same position, then connecting the two nucleotides in the blunt end bp to form a closed hairpin at each DNA site^12^. After the first strand is nicked, there is an intermediate conformational state referred to as the hairpin-forming complex (HFC), in which the active site is formed to mediate cleavage of the second strand and then form the hairpin. In the HFC structure (PDB: 5ZE1), two Mn^2+^ ions form inner sphere interactions at the active site with catalytic amino acids as well as the backbone of the two nucleotides at position *n*-1, thus facilitating the formation of the DNA connection^13^. The A-T bp at position *n*-2 in this structure has weak electron density and was modeled as Watson-Crick bp (Extended Data Fig. 11).

Interestingly, in a catalytically inactive HFC structure (PDB: 5ZDZ) bound to Ca^2+^, which is inhibitory to catalysis, the active site is not fully formed. In this structure, only one Ca^2+^ is bound and forms a coordination distinct from that found in the active state of the enzyme^13^ (Extended Data Fig. 11). Interestingly, in this structure, an ambiguous A-T bp that is slightly more favored as Hoogsteen is observed at position *n*-2 (Extended Data Fig. 7 and 11). The newly positioned Ca^2+^ appears to induce a conformational change at the *n*-1 thymine through inner sphere coordination with the thymine phosphate, and a Hoogsteen bp is observed at the neighboring position *n*-2. Interestingly, in the structure of the I-DOMI-DNA complex, a change in Mn^2+^ coordination also leads to a conformational change at position *n*-1, and a Hoogsteen bp is observed at position *n*-2.

**Supplementary Note 6: Ambiguous Hoogsteen bps in DNA terminal ends**

In addition to the terminal Hoogsteen bps found in the DNA sites bound to the homing endonuclease I-Onul (PDB: 3QQY) and the regulatory protein Esp1396I (PDB: 4IWR), our analysis also identified an additional seven A-T ambiguous Hoogsteen bps at DNA terminal ends (Supplementary Table 7). Four of these bps are tandem A-T (A1-T16 and A15-T2) bps located at the DNA terminal ends in two crystal structures (PDB: 6FQP, 6FQQ) of transcription factor TGIF1 bound to a duplex DNA (Extended Data Fig. 7 and Supplementary Table 7). Although these Hoogsteen bps are within the protein binding site and form van der Waals contacts with Arg167 in the wild-type complex (PDB: 6FQP), they are also seen in the absence of these contacts in structures of the TGIF1(R167A-R168A)-DNA (PDB: 6FQQ). This indicates that the Hoogsteen bps are unlikely to arise from protein-DNA interactions. The remaining three terminal Hoogsteen bps in protein-DNA complexes are distant from the protein binding site and include the Glucocorticoid receptor-DNA (PDB: 5E6C), Max-DNA (PDB: 5EYO) and Capicua HMG-box domain-DNA (PDB: 6JRP) (Extended Data Fig. 7 and Supplementary Table 7). All of these Hoogsteen bps appear to be stabilized by contacts from symmetry related molecules in the crystals.

**Extended Data Figures**

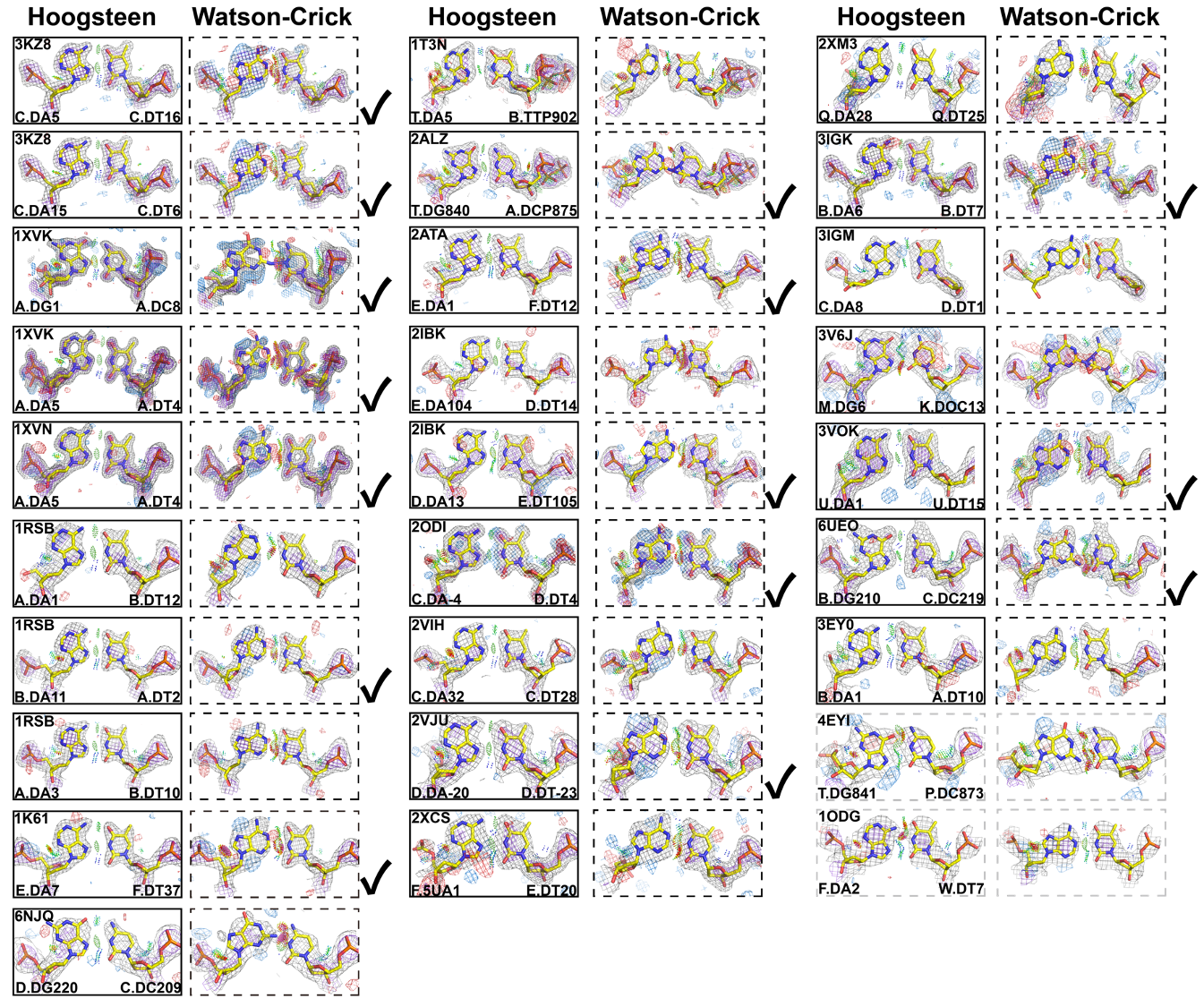

**Extended Data Fig. 1 The Hoogsteen and Watson-Crick models in the *training* dataset.** Comparison of 2mF_o_-DF_c_ and mF_o_-DF_c_ electron density maps with the original Hoogsteen model (left, solid boxes) and corresponding mismodeled Watson-Crick model (right, dashed boxes) for bps used in the *training* dataset (Supplementary Table 1). Electron density meshes and stereochemistry are as described in Fig. 1c and the box scheme is as described in Fig. 2a. Check marks at the right of the boxes indicate that the mismodeled Watson-Crick can also be identified by the *find_purine_decoy* program in Hintze *et al*. The electron density for the bps in a Vsr enzyme DNA complex (PDB: 1ODG) and in a human polymerase ι DNA complex (PDB: 4EYI) were too poor to permit discrimination between a Watson-Crick versus a Hoogsteen bp, though in the Vsr enzyme DNA, modeling Watson-Crick results in slightly better stereochemistry than modeling the bp with a Hoogsteen.

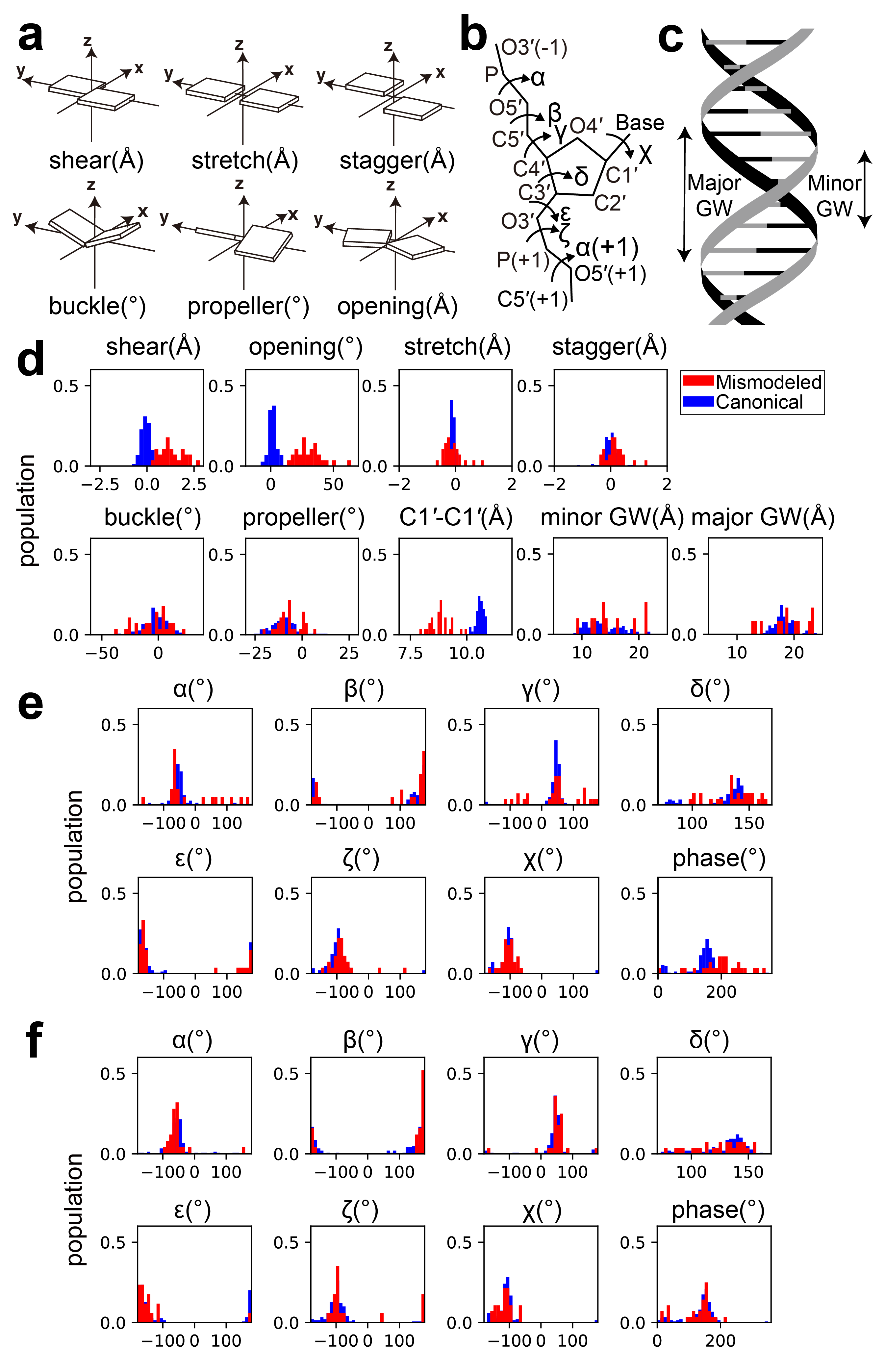

**Extended Data Fig. 2 Structural fingerprints of mismodeled Watson-Crick base pairs.** **(a-c)** Definition of **(a)** bp parameters, **(b)** local torsion angles and **(c)** major and minor groove widths. **(d-f)** 1D histograms showing **(d)** bp parameters, C1′-C1′ distance and major/minor groove widths **(e)** purine torsion angles **(f)** pyrimidine torsion angles for mismodeled Watson-Crick bps (red) versus canonical Watson-Crick bps (blue) for the *training* dataset.

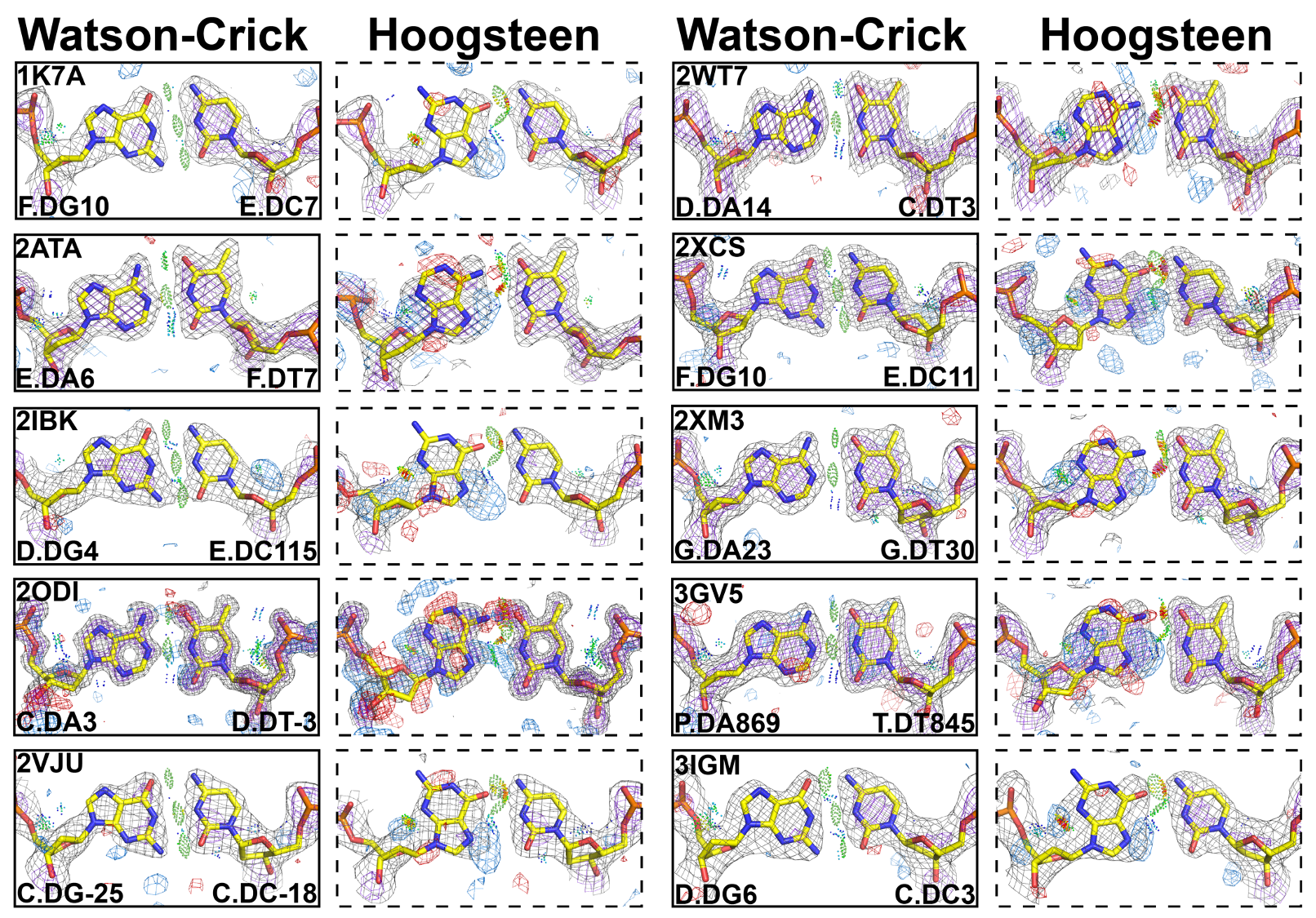

**Extended Data Fig. 3 The Hoogsteen and Watson-Crick models in *negative training* dataset.** Comparison of 2mF_o_-DF_c_ and mF_o_-DF_c_ electron density maps with the original Watson-Crick model (left, solid skyblue boxes) and corresponding mismodeled Hoogsteen bp model (right, dashed boxes) for the bps in the *negative training* dataset (Supplementary Table 2). Electron density meshes and stereochemistry are as described in Fig. 1c and the box scheme is as described in Fig. 2a.

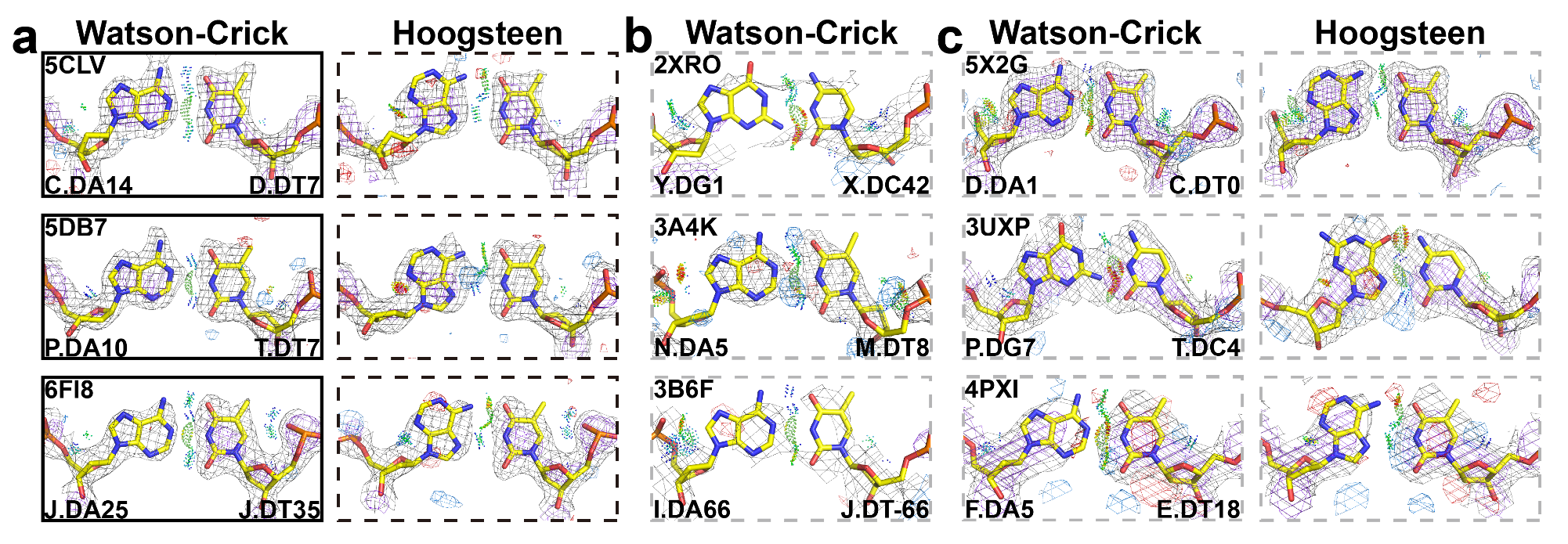

**Extended Data Fig. 4 Representative examples of Watson-Crick base pairs and ambiguous base pairs identified in the structure-based screen.** Shown are 2mF_o_-DF_c_ and mF_o_-DF_c_ electron density maps for **(a)** bps that are better modeled as Watson-Crick (left, solid skyblue boxes) relative to Hoogsteen (right, dashed boxes), **(b)** ambiguous bps with weak local density (dashed boxes) **(c)** ambiguous bps showing similar agreement with the Watson-Crick (left, dashed boxes) and Hoogsteen models (right, dashed boxes). Electron density meshes and stereochemistry are as described in Fig. 1c and the box scheme is as described in Fig. 2a. The dataset from the structure-based screen is summarized in Supplementary Table 3, 4.

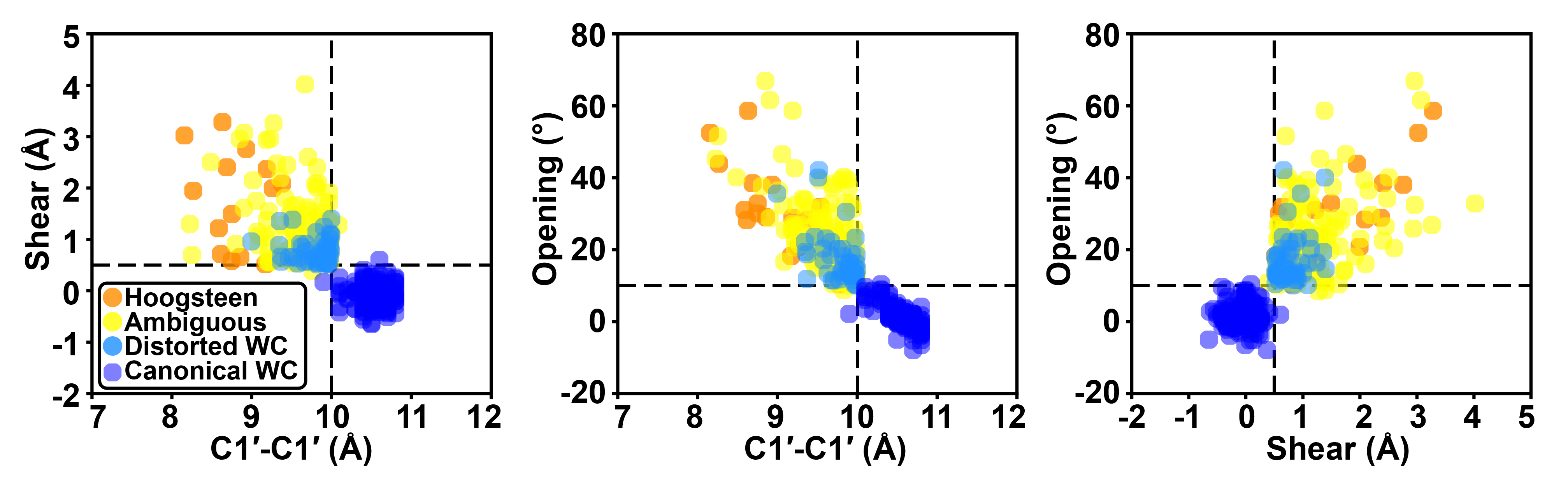

**Extended Data Fig. 5 2D scatter plot showing base pair parameters for base pairs identified using the structure-based screening (after data curation).** Hoogsteen, ambiguous, distorted Watson-Crick (Distorted WC) bps from the structure-based screening and the canonical Watson-Crick (Canonical WC) bps from Zhou *et al.* are colored in orange, yellow, skyblue, and blue, respectively. The three structural criteria used in the structure-based screening are denoted as the dashed line. The distorted Watson-Crick bps satisfying all three structural criteria are enriched at the edge of the structural criteria cutoff.

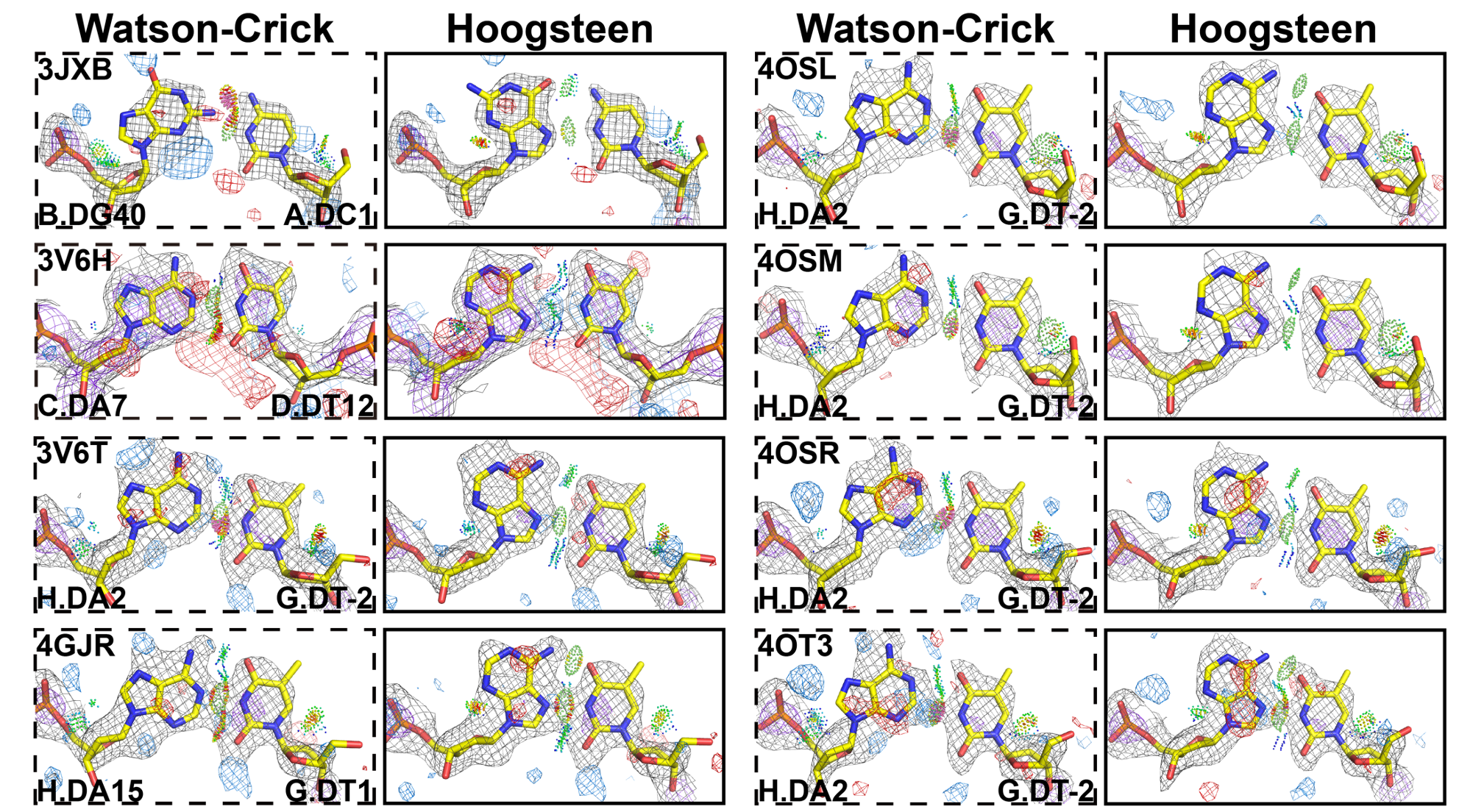

**Extended Data Fig. 6 Additional examples of Hoogsteen base pairs identified in the study.** Comparison of 2mF_o_-DF_c_ and mF_o_-DF_c_ electron density maps for the original Watson-Crick (left, dashed boxes) and the corresponding Hoogsteen models (right, solid orange boxes). Electron density meshes and stereochemistry are as described in Fig. 1c and the box scheme is as described in Fig. 2a. The entire dataset is summarized in Supplementary Table 6.

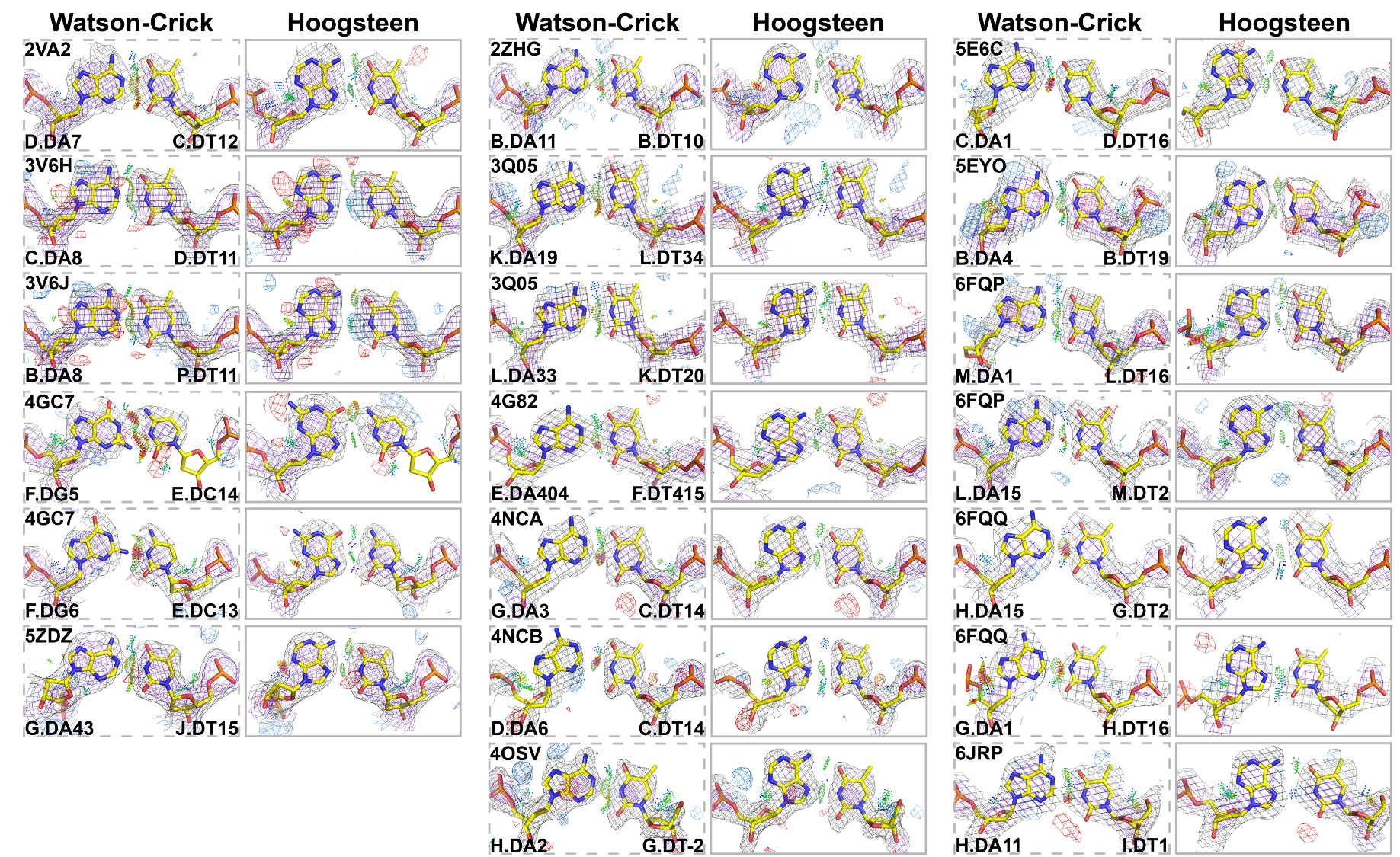

**Extended Data Fig. 7 Additional examples of ambiguous Hoogsteen base pairs.** Comparison of 2mF_o_-DF_c_ and mF_o_-DF_c_ electron density maps for the original Watson-Crick (left, dashed boxes) and the corresponding Hoogsteen models (right, solid yellow boxes). Electron density meshes and stereochemistry are as described in Fig. 1c and the box scheme is as described in Fig. 2a. The entire dataset is summarized in Supplementary Table 7.

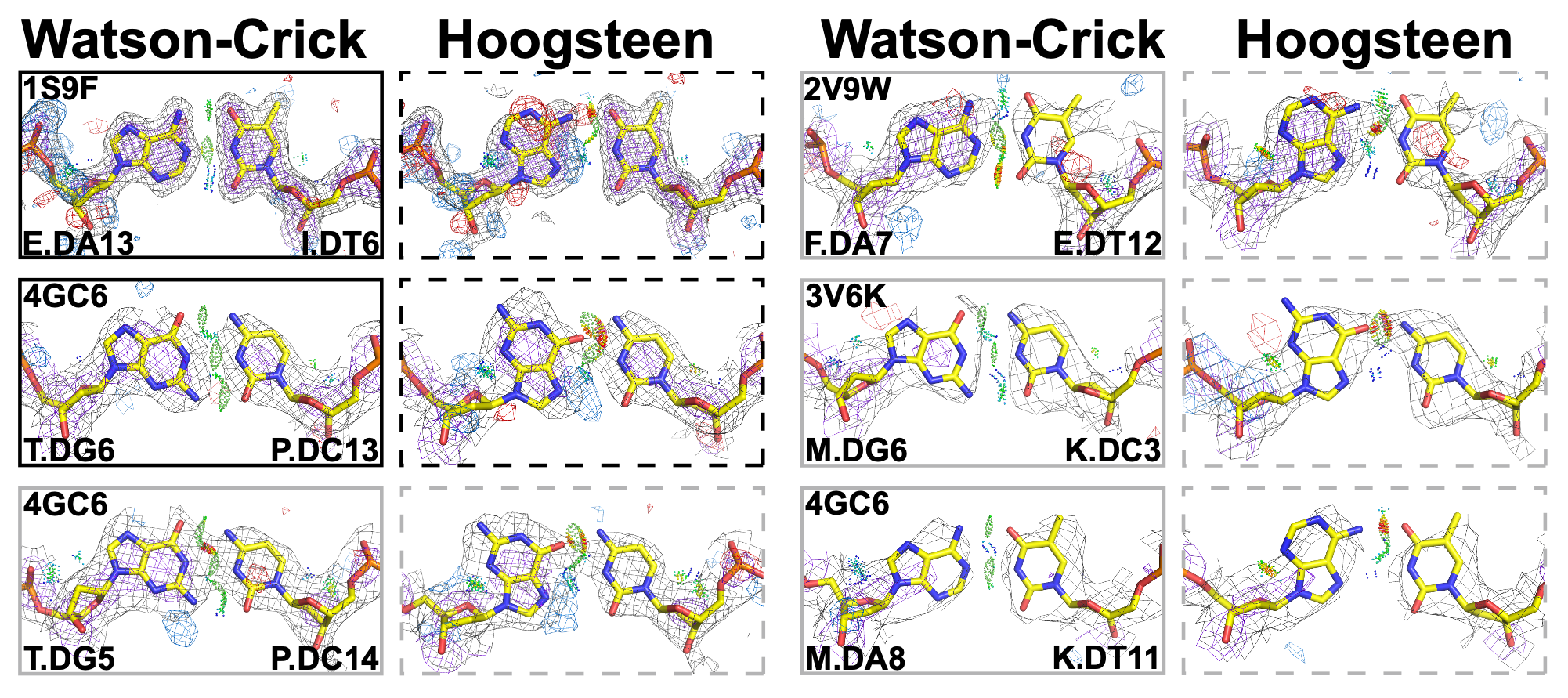

**Extended Data Fig. 8 Additional examples of Watson-Crick or ambiguous Watson-Crick base pairs in Dpo4-DNA.** Comparison of 2mF_o_-DF_c_ and mF_o_-DF_c_ electron density maps for the original Watson-Crick (left, solid boxes) and the corresponding Hoogsteen models (right, dashed boxes). The boxes of Watson-Crick model are colored as skyblue and yellow for Watson-Crick and ambiguous Watson-Crick, respectively. Electron density meshes and stereochemistry are as described in Fig. 1c and the box scheme is as described in Fig. 2a. The corresponding schematic structures are shown in Fig. 2.

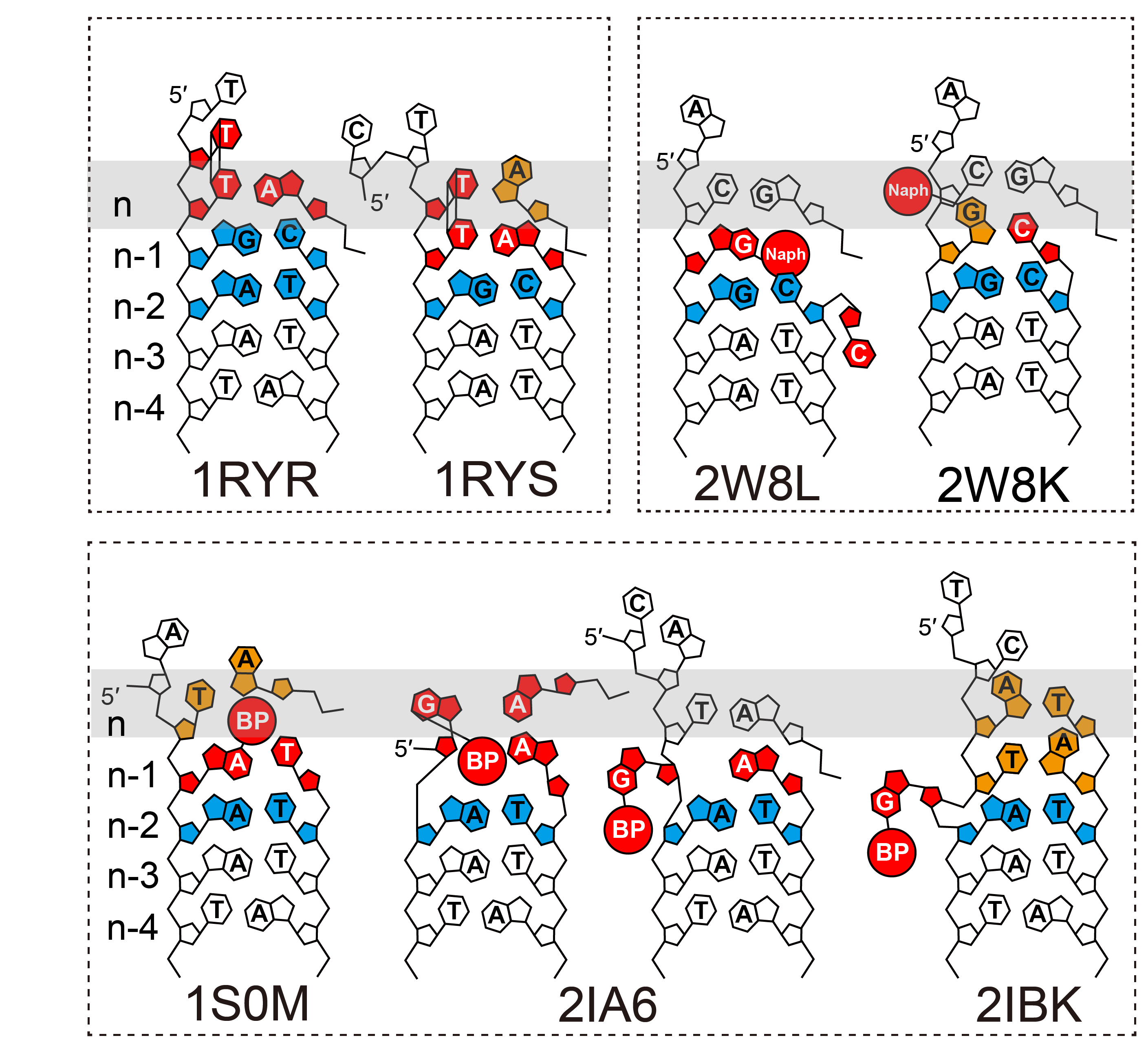

**Extended Data Fig. 9 Schematic structures of previously reported Hoogsteen bps in Dpo4.** Schematic showing the DNA containing Hoogsteen (in orange) and Watson-Crick (in skyblue) bps as well as the lesions (in red). DNA structures with similar lesions are in the same dashed box. BP = Benzo[a]pyrene adduct, Naph = Naphthyl adduct. For PDB 2IA6, the two different complexes in the same ASU reveal different DNA conformations.

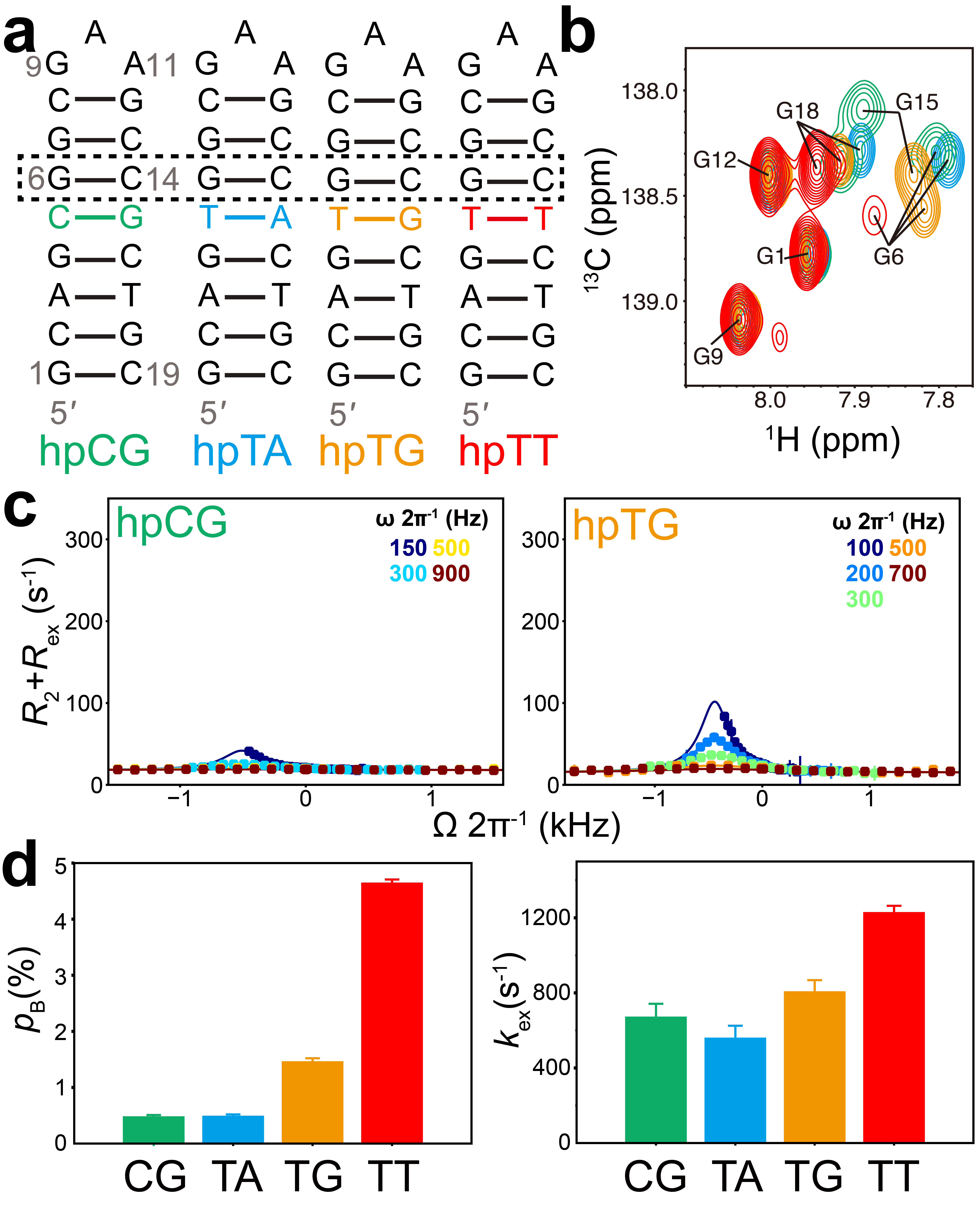

**Extended Data Fig. 10 NMR reveals enhanced propensities to form Hoogsteen bp when flanking mismatches.** **(a)** Structure of the hairpin DNA construct with and without the mismatch (in color) highlighting the G6-C14 bp (dashed rectangle) targeted for NMR relaxation dispersion measurements. **(b)** 2D [^13^C, ^1^H] HSQC of DNA aromatic region of the various constructs (color coded). **(c)** Off-resonance *R*_1ρ_ profiles for G6-C8 in hpCG and hpTG. Spin-lock powers are color coded. Error bars were estimated using a Monte-Carlo scheme and are smaller than data points (Methods). **(d)** Bar plot of p_B_ and *k*_ex_ of NMR *R*_1ρ_ profiles. Error bars denote the *R*_1ρ_ fitting errors.

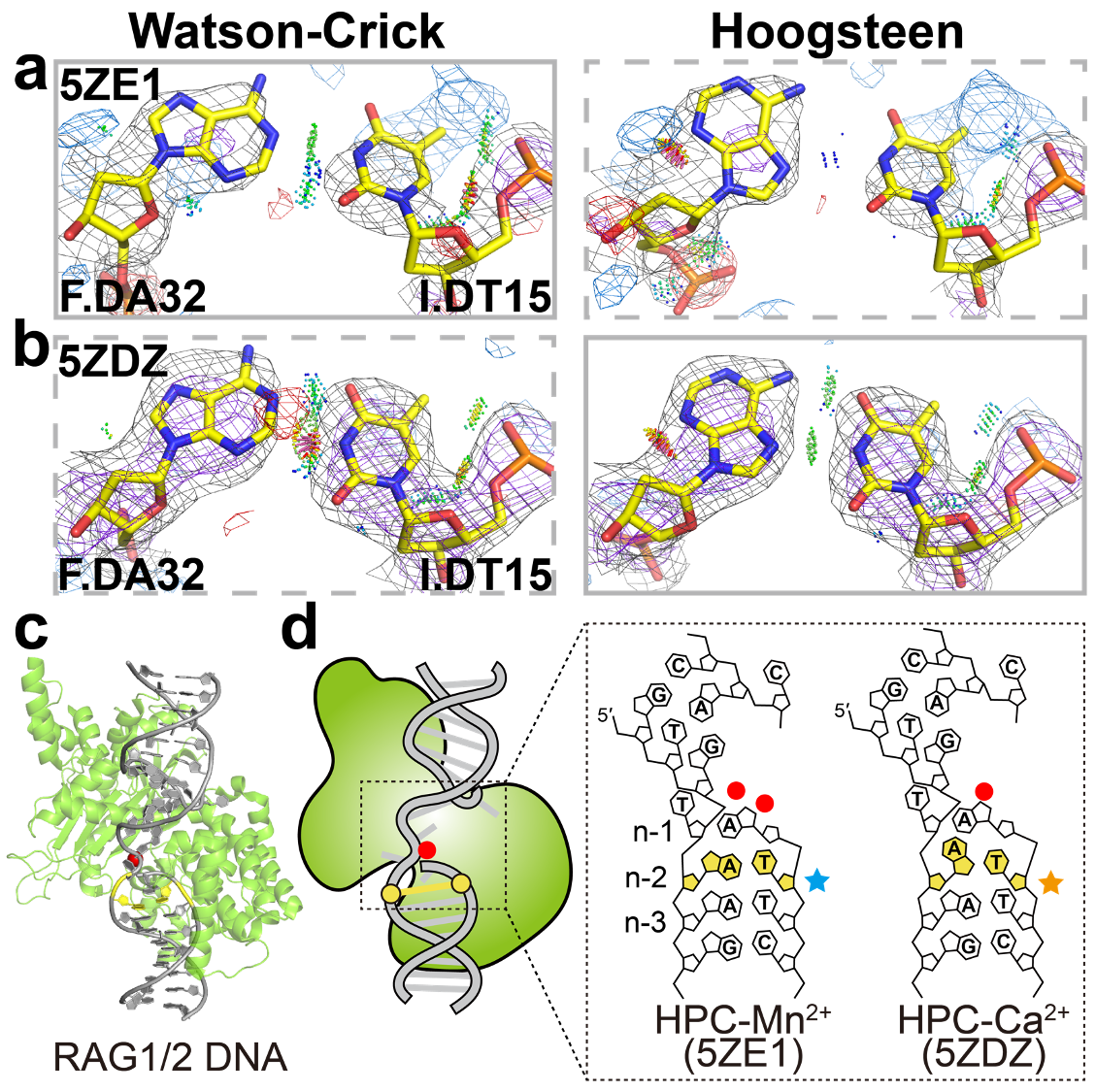

**Extended Data Fig. 11 Hoogsteen base pairs in recombinase RAG1/2. (a,b)** Comparison of 2mF_o_-DF_c_ and mF_o_-DF_c_ electron density maps for the original Watson-Crick (left) and the corresponding Hoogsteen models (right) for the A-T bp at position *n*-2 in the hairpin forming complex structure bound with **(a)** Mn^2+^ or **(b)** Ca^2+^ in the active site. Electron density meshes and stereochemistry are as described in Fig. 1c and the box scheme is as described in Fig. 2a. **(c)** 3D structures of the protein-DNA complex showing the ambiguous bp. **(d)** Schematic showing the DNA containing ambiguous bps (in yellow). Also shown are the metal ions (red filled circle).

**Supplementary Tables**

**Supplementary Table 1. Base pair entries in the *training* dataset**

| PDB ID | Resolution  (Å)^a^ | Bp | nt_1^b^ | nt_2 | WC  clash^c^ | Shear  (Å) | Opening  (°) | C1′-C1′  (Å) | Note |
| --- | --- | --- | --- | --- | --- | --- | --- | --- | --- |
| 1K61 | 2.1 | AT | E.DA7 | F.DT37 | TRUE | 1.548 | 33.98 | 8.726 |  |
| 1ODG | 2.8 | AT | F.DA2 | W.DT7 | FALSE | 0.467 | 14.248 | 9.94 | ambiguous |
| 1RSB | 2.17 | AT | A.DA1 | B.DT12 | TRUE | 2.639 | 45.089 | 8.381 |  |
| 1RSB | 2.17 | AT | A.DT2 | B.DA11 | TRUE | 1.697 | 31.365 | 8.876 |  |
| 1RSB | 2.17 | AT | A.DA3 | B.DT10 | TRUE | 1.292 | 29.232 | 8.836 |  |
| 1T3N | 2.3 | At | B.TTP902 | T.DA5 | TRUE | 2.289 | 36.522 | 8.769 | weak *omit* map |
| 1XVK | 1.26 | CG | A.DG1 | A.DC8 | TRUE | 1.833 | 42.502 | 8.867 |  |
| 1XVK | 1.26 | AT | A.DT4 | A.DA5 | TRUE | 2.026 | 35.258 | 8.424 |  |
| 1XVN | 1.5 | AT | A.DT4 | A.DA5 | TRUE | 1.286 | 23.373 | 8.746 |  |
| 2ALZ | 2.5 | cG | T.DG840 | A.DCP875 | TRUE | 1.298 | 32.911 | 8.862 |  |
| 2ATA | 2.2 | AT | E.DA1 | F.DT12 | TRUE | 0.981 | 21.177 | 9.003 |  |
| 2IBK | 2.25 | AT | D.DA13 | E.DT105 | TRUE | 0.375 | 35.084 | 8.426 |  |
| 2IBK | 2.25 | AT | D.DT14 | E.DA104 | TRUE | 1.027 | 37.164 | 8.159 |  |
| 2ODI | 1.45 | AT | C.DA-4 | D.DT4 | TRUE | 1.884 | 26.596 | 9.032 |  |
| 2VIH | 2.1 | AT | C.DT28 | C.DA32 | TRUE | 2.116 | 39.932 | 8.542 |  |
| 2VJU | 2.4 | AT | D.DT-23 | D.DA-20 | TRUE | 1.076 | 22.957 | 9.068 |  |
| 2XCS | 2.1 | AT | E.DT20 | F.DA1 | TRUE | 0.823 | 28.115 | 8.36 |  |
| 2XM3 | 2.3 | AT | Q.DT25 | Q.DA28 | TRUE | 1.885 | 36.135 | 8.462 |  |
| 3EY0 | 2.52 | AT | A.DT10 | B.DA1 | FALSE | 0.684 | 19.656 | 9.322 |  |
| 3IGK | 1.7 | AT | B.DA6 | B.DT7 | TRUE | 1.103 | 25.419 | 8.887 |  |
| 3IGM | 2.2 | AT | C.DA8 | D.DT1 | TRUE | 1.426 | 16.739 | 9.378 |  |
| 3KZ8 | 1.91 | AT | C.DA5 | C.DT16 | TRUE | 1.189 | 25.545 | 8.762 |  |
| 3KZ8 | 1.91 | AT | C.DT6 | C.DA15 | TRUE | 1.129 | 25.241 | 8.832 |  |
| 3V6J | 2.3 | cG | K.DOC13 | M.DG6 | TRUE | 0.522 | 51.906 | 7.911 |  |
| 3VOK | 2 | AT | U.DA1 | U.DT15 | FALSE | 0.793 | 20.137 | 9.217 |  |
| 4EYI | 2.9 | CG | T.DG841 | P.DC873 | TRUE | 0.9 | 17.619 | 9.7 | ambiguous |
| 6NJQ | 2.75 | CG | C.DC209 | D.DG220 | TRUE | 2.285 | 61.048 | 8.222 |  |
| 6UEO | 2 | CG | B.DG210 | C.DC219 | TRUE | 0.551 | 26.664 | 9.378 |  |

a. The resolutions are obtained directly from the RCSB PDB.

b. Nucleotide identifier: e.g. X.DA1 = adenine 1 in chain X

c. Stereo clash between the two bases in the Watson-Crick model

**Supplementary Table 2. Base pair entries in the *negative* *training* dataset**

| PDB ID | Resolution  (Å) | Bp | nt_1 | nt_2 | WC  clash | Shear  (Å) | Opening  (°) | C1′-C1′  (Å) | Note |
| --- | --- | --- | --- | --- | --- | --- | --- | --- | --- |
| 1K7A | 2.8 | CG | E.DC7 | F.DG10 | FALSE | -0.299 | 0.568 | 10.51 |  |
| 2ATA | 2.2 | AT | E.DA6 | F.DT7 | FALSE | -0.018 | 7.468 | 10.32 |  |
| 2IBK | 2.25 | CG | D.DG4 | E.DC115 | FALSE | 0.028 | 1.884 | 10.62 |  |
| 2ODI | 1.45 | AT | C.DA3 | D.DT-3 | FALSE | 0.273 | 0.245 | 10.488 |  |
| 2VJU | 2.4 | CG | C.DG-25 | C.DC-18 | FALSE | -0.308 | 1.784 | 10.461 |  |
| 2WT7 | 2.3 | AT | C.DT3 | D.DA14 | FALSE | 0.006 | 7.856 | 10.397 |  |
| 2XCS | 2.1 | CG | E.DC11 | F.DG10 | FALSE | -0.165 | -0.226 | 10.509 |  |
| 2XM3 | 2.3 | AT | G.DA23 | G.DT30 | FALSE | 0.074 | 0.06 | 10.539 |  |
| 3GV5 | 2 | AT | P.DA869 | T.DT845 | FALSE | 0.099 | -0.648 | 10.61 |  |
| 3IGM | 2.2 | CG | C.DC3 | D.DG6 | FALSE | 0.132 | -5.154 | 10.784 |  |

**Supplementary Table 3. Statistics for base pair entries in the structure-based survey**

| Dataset | Number of bps  (Before curation) | Number of bps  (After curation) |
| --- | --- | --- |
| *Parent* | 97,100 | N/A |
| *Starting* | 215 | N/A |
| *Filtered* | 66 | N/A |
| *Hoogsteen* | 22 | 17 |
| *Watson-Crick* | 58 | 52 |
| *Ambiguous* | 135 | 130 |

**Supplementary Table 4. All base pair entries from the structure-based screening in this study^a^**

| PDB ID | Resolution (Å) | Bp | nt_1 | nt_2 | Result^b^ | Context^c^ | Primary^d^ | Stress^e^ | WC_clash | Note^f^ |
| --- | --- | --- | --- | --- | --- | --- | --- | --- | --- | --- |
| 1S97 | 2.4 | AT | E.DT12 | I.DA7 | HG | Pol_n-2 | TRUE | TRUE | TRUE | Hintze *et al.* |
| 1S97 | 2.4 | AT | F.DT12 | J.DA7 | HG | Pol_n-2 | FALSE | TRUE | TRUE | Hintze *et al.* |
| 1S97 | 2.4 | AT | G.DT12 | K.DA7 | HG | Pol_n-2 | FALSE | TRUE | TRUE | Hintze *et al.* |
| 1S97 | 2.4 | AT | H.DT12 | L.DA7 | HG | Pol_n-2 | FALSE | TRUE | TRUE | Hintze *et al.* |
| 3JXB | 1.67 | CG | A.DC1 | B.DG40 | HG | Terminal | TRUE | TRUE | TRUE | Hintze *et al.* |
| 3QQY | 2.401 | CG | B.DC25 | C.DG1 | HG | Terminal | TRUE | TRUE | TRUE |  |
| 3V6H | 2.3 | AT | C.DA7 | D.DT12 | HG | Pol_n-2 | TRUE | TRUE | TRUE |  |
| 3V6H | 2.3 | AT | T.DA7 | P.DT12 | HG | Pol_n-2 | FALSE | TRUE | TRUE |  |
| 3V6J | 2.3 | AT | P.DT12 | B.DA7 | HG | Pol_n-2 | TRUE | TRUE | TRUE | Hintze *et al.* |
| 3V6J | 2.3 | AT | K.DT12 | M.DA7 | HG | Pol_n-2 | FALSE | TRUE | TRUE | Hintze *et al.* |
| 3V6T | 1.85 | AT | G.DT-2 | H.DA2 | HG | Terminal | TRUE | TRUE | TRUE | Hintze *et al.* |
| 4GJR | 1.85 | AT | G.DT1 | H.DA15 | HG | Terminal | TRUE | TRUE | TRUE |  |
| 4IWR | 2.4 | AT | C.DA1 | D.DT25 | HG | Terminal | TRUE | TRUE | TRUE |  |
| 4IWR | 2.4 | AT | G.DA1 | H.DT25 | HG | Terminal | FALSE | TRUE | TRUE |  |
| 4KPY | 2.406 | AT | C.DT14 | N.DA6 | HG | Helical | TRUE | FALSE | FALSE |  |
| 4KPY | 2.406 | AT | E.DT14 | M.DA6 | HG | Helical | FALSE | FALSE | TRUE |  |
| 4OSL | 2.447 | AT | G.DT-2 | H.DA2 | HG | Terminal | TRUE | TRUE | TRUE |  |
| 4OSM | 2.454 | AT | G.DT-2 | H.DA2 | HG | Terminal | TRUE | TRUE | TRUE |  |
| 4OSR | 1.944 | AT | G.DT-2 | H.DA2 | HG | Terminal | TRUE | TRUE | TRUE |  |
| 4OT3 | 1.944 | AT | G.DT-2 | H.DA2 | HG | Terminal | TRUE | TRUE | TRUE |  |
| 5A0W | 2.2 | CG | B.DG11 | C.DC15 | HG | Helical | TRUE | TRUE | TRUE |  |
| 5A0W | 2.2 | CG | E.DG11 | F.DC15 | HG | Helical | FALSE | TRUE | TRUE |  |
| 5A0W | 2.2 | CG | H.DG11 | I.DC15 | HG | Helical | FALSE | TRUE | TRUE |  |
| 5HP4 | 1.86 | AT | X.DT5 | X.DA6 | HG | Helical | TRUE | TRUE | TRUE |  |
| 5WN0 | 2.6 | AT | C.DT1 | E.DA11 | HG | Nick | TRUE | TRUE | TRUE |  |
| 5WN0 | 2.6 | CG | C.DC2 | E.DG10 | HG | Nick-1 | TRUE | TRUE | TRUE |  |
| 1HCR | 2.3 | AT | B.DA13 | C.DT19 | HG=WC | Helical | TRUE | FALSE | FALSE | Remove Lys187 |
| 2VA2 | 2.8 | AT | E.DT12 | F.DA7 | HG=WC | Pol_n-2 | FALSE | TRUE | FALSE |  |
| 3Q05 | 2.4 | AT | K.DA9 | L.DT44 | HG=WC | Helical | FALSE | FALSE | FALSE |  |
| 3TS8 | 2.8 | AT | K.DA19 | L.DT34 | HG=WC | Helical | TRUE | FALSE | TRUE |  |
| 3UXP | 2.723 | CG | P.DG7 | T.DC4 | HG=WC | Pol_n-1 | TRUE | TRUE | TRUE |  |
| 4G82 | 3.1 | AT | E.DT405 | F.DA414 | HG=WC | Helical | FALSE | FALSE | TRUE |  |
| 4IVZ | 3.1 | AT | C.DA1 | H.DT1 | HG=WC | Sticky_artifact | TRUE | TRUE | TRUE |  |
| 4NNU | 2.81 | AT | C.DT12 | D.DA11 | HG=WC | Helical | TRUE | TRUE | TRUE |  |
| 4PXI | 3.2 | AT | E.DT18 | F.DA5 | HG=WC | Helical | TRUE | FALSE | FALSE |  |
| 5K1Y | 2.97 | AT | P.DA21 | N.DT30 | HG=WC | Helical | TRUE | FALSE | FALSE |  |
| 5NFV | 2.501 | AT | C.DA-25 | D.DT25 | HG=WC | Helical | TRUE | TRUE | TRUE |  |
| 5VFX | 2.81 | AT | I.DA1 | J.DT23 | HG=WC | Terminal | TRUE | TRUE | TRUE |  |
| 5X2G | 2.4 | AT | C.DT0 | D.DA1 | HG=WC | Terminal | TRUE | TRUE | FALSE |  |
| 5X2H | 2.3 | AT | C.DT0 | D.DA1 | HG=WC | Terminal | TRUE | TRUE | FALSE |  |
| 6DSX | 1.99 | AT | P.DA10 | T.DT5 | HG=WC | Pol_n-1 | TRUE | TRUE | TRUE | Remove Arg629 |
| 6FQP | 2.42 | AT | L.DA14 | M.DT3 | HG=WC | Helical | TRUE | FALSE | FALSE |  |
| 6FQP | 2.42 | AT | L.DA1 | M.DT16 | HG=WC | Terminal | FALSE | TRUE | TRUE |  |
| 6FQP | 2.42 | AT | L.DT2 | M.DA15 | HG=WC | Terminal-1 | FALSE | FALSE | FALSE |  |
| 6G1L | 2.4 | CG | B.DG2 | B.DC15 | HG=WC | Terminal-1 | TRUE | TRUE | TRUE |  |
| 2VA2 | 2.8 | AT | C.DT12 | D.DA7 | HG>WC | Pol_n-2 | TRUE | TRUE | TRUE |  |
| 2ZHG | 2.8 | AT | B.DT10 | B.DA11 | HG>WC | Helical | TRUE | FALSE | TRUE |  |
| 3Q05 | 2.4 | AT | K.DA19 | L.DT34 | HG>WC | Helical | TRUE | FALSE | TRUE |  |
| 3Q05 | 2.4 | AT | K.DT20 | L.DA33 | HG>WC | Helical | TRUE | FALSE | FALSE |  |
| 3V6H | 2.3 | AT | C.DA8 | D.DT11 | HG>WC | Helical | TRUE | FALSE | FALSE |  |
| 3V6H | 2.3 | AT | T.DA8 | P.DT11 | HG>WC | Helical | FALSE | FALSE | FALSE |  |
| 3V6J | 2.3 | AT | P.DT11 | B.DA8 | HG>WC | Helical | TRUE | FALSE | TRUE |  |
| 3V6J | 2.3 | AT | K.DT11 | M.DA8 | HG>WC | Helical | FALSE | FALSE | TRUE |  |
| 4G82 | 3.1 | AT | E.DA404 | F.DT415 | HG>WC | Helical | TRUE | FALSE | TRUE |  |
| 4GC7 | 2.89 | CG | E.DC13 | F.DG6 | HG>WC | Pol_n-2 | TRUE | FALSE | TRUE |  |
| 4GC7 | 2.89 | CG | E.DC14 | F.DG5 | HG>WC | Pol_n-1 | TRUE | TRUE | TRUE |  |
| 4GC7 | 2.89 | CG | C.DC13 | D.DG6 | HG>WC | Pol_n-2 | FALSE | FALSE | TRUE |  |
| 4GC7 | 2.89 | CG | C.DC14 | D.DG5 | HG>WC | Pol_n-1 | FALSE | TRUE | TRUE |  |
| 4NCA | 2.489 | AT | C.DT14 | G.DA3 | HG>WC | Helical | TRUE | FALSE | TRUE |  |
| 4NCA | 2.489 | AT | E.DT14 | H.DA6 | HG>WC | Helical | FALSE | FALSE | TRUE |  |
| 4NCB | 2.189 | AT | C.DT14 | D.DA6 | HG>WC | Terminal-1 | TRUE | TRUE | TRUE |  |
| 4NCB | 2.189 | AT | E.DT14 | H.DA6 | HG>WC | Helical | FALSE | FALSE | TRUE |  |
| 4OSV | 1.996 | AT | G.DT-2 | H.DA2 | HG>WC | Terminal | TRUE | TRUE | FALSE |  |
| 5E6C | 2.2 | AT | C.DA1 | D.DT16 | HG>WC | Terminal | TRUE | TRUE | TRUE |  |
| 5EYO | 2.39 | AT | B.DA4 | B.DT19 | HG>WC | Terminal | TRUE | TRUE | FALSE |  |
| 5EYO | 2.39 | AT | D.DA4 | D.DT19 | HG>WC | Terminal | FALSE | TRUE | FALSE |  |
| 5ZDZ | 2.8 | AT | F.DA32 | I.DT15 | HG>WC | Nick-1 | TRUE | TRUE | TRUE |  |
| 5ZDZ | 2.8 | AT | J.DT15 | G.DA43 | HG>WC | Nick-1 | TRUE | TRUE | TRUE |  |
| 6FQP | 2.42 | AT | L.DA15 | M.DT2 | HG>WC | Terminal-1 | TRUE | FALSE | TRUE |  |
| 6FQP | 2.42 | AT | L.DT16 | M.DA1 | HG>WC | Terminal | TRUE | TRUE | FALSE |  |
| 6FQQ | 3.25 | AT | G.DA1 | H.DT16 | HG>WC | Terminal | TRUE | TRUE | TRUE |  |
| 6FQQ | 3.25 | AT | G.DT2 | H.DA15 | HG>WC | Terminal-1 | TRUE | TRUE | TRUE |  |
| 6FQQ | 3.25 | AT | L.DA1 | M.DT16 | HG>WC | Terminal | FALSE | TRUE | TRUE |  |
| 6FQQ | 3.25 | AT | L.DT2 | M.DA15 | HG>WC | Terminal-1 | FALSE | FALSE | TRUE |  |
| 6JRP | 3 | AT | H.DA11 | I.DT1 | HG>WC | Terminal | TRUE | TRUE | TRUE |  |
| 1F4K | 2.5 | AT | D.DA11 | E.DT11 | Low | Helical | TRUE | FALSE | FALSE |  |
| 1RUO | 2.7 | AT | C.DT6 | F.DA6 | Low | Helical | TRUE | FALSE | FALSE |  |
| 2ASJ | 2.35 | AT | D.DA807 | E.DT913 | Low | Helical | TRUE | FALSE | TRUE |  |
| 2DY4 | 2.65 | CG | K.DG16 | L.DC102 | Low | Terminal-1 | TRUE | TRUE | TRUE |  |
| 2HHQ | 1.8 | CG | B.DC22 | C.DG16 | Low | Terminal | TRUE | TRUE | TRUE |  |
| 2P6R | 3 | AT | X.DA1 | Y.DT25 | Low | Terminal | TRUE | TRUE | TRUE |  |
| 2V1U | 3.1 | AT | B.DT3 | C.DA20 | Low | Helical | TRUE | FALSE | TRUE |  |
| 2XHI | 1.55 | CG | B.DC15 | C.DG16 | Low | Terminal | TRUE | TRUE | TRUE |  |
| 2XRO | 3.4 | CG | X.DC42 | Y.DG1 | Low | Terminal | TRUE | TRUE | TRUE |  |
| 3A4K | 2.17 | AT | M.DT8 | N.DA5 | Low | Helical | TRUE | FALSE | FALSE |  |
| 3B6F | 3.45 | AT | I.DA66 | J.DT-66 | Low | Helical | TRUE | TRUE | FALSE |  |
| 3JR4 | 2.601 | AT | B.DA5 | C.DT13 | Low | Terminal | TRUE | TRUE | FALSE |  |
| 3KMP | 2.7 | AT | C.DT2 | D.DA16 | Low | Terminal | TRUE | TRUE | TRUE |  |
| 3KTU | 2.3 | AT | B.DA12 | C.DT19 | Low | Terminal-1 | TRUE | TRUE | TRUE |  |
| 3KUY | 2.9 | AT | I.DA59 | J.DT-59 | Low | Helical | TRUE | FALSE | FALSE |  |
| 3L2Q | 3.25 | AT | C.DA19 | D.DT1 | Low | Terminal | TRUE | TRUE | FALSE |  |
| 3L2R | 2.88 | AT | C.DA19 | D.DT1 | Low | Terminal | TRUE | TRUE | FALSE |  |
| 3L2U | 3.15 | AT | C.DA19 | D.DT1 | Low | Terminal | TRUE | TRUE | FALSE |  |
| 3LZ1 | 2.5 | AT | I.DA-68 | J.DT68 | Low | Helical | TRUE | FALSE | TRUE |  |
| 3MGP | 2.44 | AT | I.DA-73 | J.DT73 | Low | Terminal | TRUE | TRUE | FALSE |  |
| 3MGV | 2.29 | AT | F.DT5 | H.DA2 | Low | Nick-1 | TRUE | FALSE | FALSE |  |
| 3PIH | 2.9 | AT | D.DA1 | D.DT32 | Low | Terminal | TRUE | TRUE | TRUE |  |
| 3PW7 | 2.9 | AT | F.DT377 | G.DA356 | Low | Helical | TRUE | FALSE | FALSE |  |
| 3Q06 | 3.2 | AT | K.DT26 | L.DA27 | Low | Terminal | TRUE | TRUE | FALSE |  |
| 3S5A | 1.7 | AT | B.DT260 | C.DA284 | Low | Terminal | TRUE | TRUE | FALSE |  |
| 3U61 | 3.2 | AT | I.DA8 | J.DT3 | Low | Terminal | TRUE | TRUE | TRUE |  |
| 3V6J | 2.3 | CG | P.DG2 | B.DC17 | Low | Terminal | TRUE | TRUE | FALSE |  |
| 3V6T | 1.85 | AT | G.DT14 | H.DA-14 | Low | Terminal | TRUE | TRUE | FALSE |  |
| 4B5G | 2.75 | CG | W.DC37 | X.DG46 | Low | Flank | TRUE | TRUE | TRUE |  |
| 4B5M | 2.758 | CG | M.DC37 | X.DG46 | Low | Flank | TRUE | TRUE | TRUE |  |
| 4BXX | 3.28 | AT | N.DA5 | T.DT13 | Low | Helical | TRUE | FALSE | FALSE |  |
| 4KGC | 2.69 | AT | I.DA-18 | J.DT18 | Low | Helical | TRUE | FALSE | TRUE |  |
| 4LD9 | 3.306 | AT | I.DA-68 | J.DT68 | Low | Helical | TRUE | FALSE | FALSE |  |
| 4LD9 | 3.306 | CG | I.DC-52 | J.DG52 | Low | Helical | TRUE | TRUE | FALSE |  |
| 4LLL | 3.036 | AT | G.DT24 | H.DA1 | Low | Terminal | TRUE | TRUE | FALSE |  |
| 4MKY | 2.4 | CG | F.DG7 | H.DC10 | Low | Terminal | TRUE | TRUE | TRUE |  |
| 4NOE | 2.2 | CG | F.DG3 | F.DC6 | Low | Terminal | TRUE | TRUE | TRUE |  |
| 4OIN | 2.8 | AT | G.DA13 | H.DT15 | Low | Terminal | TRUE | TRUE | FALSE |  |
| 4OST | 1.996 | AT | G.DT-2 | H.DA2 | Low | Terminal | TRUE | TRUE | TRUE |  |
| 4OSW | 2.302 | AT | G.DT-2 | H.DA2 | Low | Terminal | TRUE | TRUE | TRUE |  |
| 4R79 | 3.1 | CG | G.DG4 | H.DC53 | Low | Terminal | TRUE | TRUE | TRUE |  |
| 4S04 | 3.2 | AT | G.DA1 | H.DT24 | Low | Terminal | TRUE | TRUE | FALSE |  |
| 4U7D | 3.4 | AT | R.DT15 | S.DA3 | Low | Terminal | TRUE | TRUE | FALSE |  |
| 4X4B | 2.8 | AT | E.DA34 | F.DT2 | Low | Terminal-1 | TRUE | FALSE | FALSE |  |
| 4X4C | 2.8 | AT | E.DA34 | F.DT2 | Low | Terminal-1 | TRUE | FALSE | FALSE |  |
| 4X4D | 2.8 | AT | E.DA34 | F.DT2 | Low | Terminal-1 | TRUE | FALSE | FALSE |  |
| 4X4E | 2.8 | AT | E.DA34 | F.DT2 | Low | Terminal-1 | TRUE | FALSE | FALSE |  |
| 4X4F | 2.8 | AT | E.DA34 | F.DT2 | Low | Terminal-1 | TRUE | FALSE | FALSE |  |
| 4X4G | 2.8 | AT | E.DA34 | F.DT2 | Low | Terminal-1 | TRUE | FALSE | FALSE |  |
| 4X4H | 2.8 | AT | E.DA34 | F.DT2 | Low | Terminal-1 | TRUE | FALSE | FALSE |  |
| 4X4I | 2.8 | AT | E.DA34 | F.DT2 | Low | Terminal-1 | TRUE | FALSE | FALSE |  |
| 4YG1 | 3.25 | AT | F.DA743 | T.DT703 | Low | Helical | TRUE | FALSE | FALSE |  |
| 4ZTU | 3.299 | CG | T.DC8 | P.DG21 | Low | Helical | TRUE | FALSE | TRUE |  |
| 4ZTZ | 3.442 | CG | T.DC8 | P.DG21 | Low | Helical | TRUE | FALSE | TRUE |  |
| 5CPI | 2.902 | AT | I.DT8 | J.DA139 | Low | Helical | TRUE | FALSE | FALSE |  |
| 5CPJ | 3.15 | AT | I.DT87 | J.DA60 | Low | Helical | TRUE | FALSE | FALSE |  |
| 5E6C | 2.2 | AT | C.DT16 | D.DA1 | Low | Terminal | TRUE | TRUE | TRUE |  |
| 5ESP | 2.995 | AT | C.DA20 | D.DT9 | Low | Helical | TRUE | FALSE | FALSE |  |
| 5F99 | 2.63 | AT | I.DT-53 | J.DA53 | Low | Helical | TRUE | FALSE | TRUE |  |
| 5GWK | 3.152 | AT | C.DA1 | F.DT20 | Low | Terminal | TRUE | TRUE | TRUE |  |
| 5GXQ | 2.85 | AT | I.DA127 | J.DT166 | Low | Helical | TRUE | FALSE | TRUE |  |
| 5OQN | 3.15 | AT | C.DA8 | D.DT11 | Low | Helical | TRUE | FALSE | FALSE |  |
| 5T9J | 3 | AT | C.DA16 | F.DT5 | Low | Terminal-1 | TRUE | FALSE | TRUE |  |
| 5TH3 | 2.33 | CG | h.DG37 | I.DC2 | Low | Terminal | TRUE | TRUE | TRUE |  |
| 5UK7 | 3 | AT | N.DA5 | M.DT16 | Low | Helical | TRUE | FALSE | TRUE |  |
| 5X6M | 3.2 | AT | G.DT19 | H.DA5 | Low | Helical | TRUE | FALSE | FALSE |  |
| 5Z30 | 2.45 | AT | I.DA1 | J.DT292 | Low | Terminal | TRUE | TRUE | FALSE |  |
| 6CG8 | 2.299 | AT | C.DT3 | F.DA17 | Low | Terminal | TRUE | TRUE | FALSE |  |
| 6DGD | 2.823 | AT | Y.DA11 | Z.DT3 | Low | Terminal | TRUE | FALSE | FALSE |  |
| 6EDC | 2.712 | AT | B.DA17 | C.DT2 | Low | Terminal | TRUE | TRUE | FALSE |  |
| 6IFM | 2.804 | AT | M.DA21 | N.DT7 | Low | Helical | TRUE | TRUE | TRUE |  |
| 6JRG | 2.005 | CG | D.DG15 | D.DC19 | Low | Terminal | TRUE | TRUE | TRUE |  |
| 6JW5 | 2.99 | AT | F.DT-2 | G.DA2 | Low | Terminal | TRUE | TRUE | FALSE |  |
| 6K1J | 2.85 | AT | I.DA-49 | J.DT49 | Low | Helical | TRUE | FALSE | FALSE |  |
| 6KDB | 2.862 | AT | E.DA433 | F.DT436 | Low | Helical | TRUE | FALSE | FALSE |  |
| 6KE9 | 2.22 | AT | I.DA29 | J.DT-29 | Low | Helical | TRUE | FALSE | FALSE |  |
| 6KE9 | 2.22 | AT | I.DT-44 | J.DA44 | Low | Helical | TRUE | FALSE | FALSE |  |
| 6KQF | 2.45 | CG | G.DC8 | H.DG20 | Low | Helical | TRUE | FALSE | TRUE |  |
| 6KQN | 3.489 | CG | G.DG12 | H.DC16 | Low | Terminal-1 | TRUE | FALSE | TRUE |  |
| 6L9H | 2.6 | AT | I.DA-55 | J.DT55 | Low | Helical | TRUE | FALSE | FALSE |  |
| 6L9H | 2.6 | AT | I.DA29 | J.DT-29 | Low | Helical | TRUE | FALSE | FALSE |  |
| 6L9H | 2.6 | AT | I.DT64 | J.DA-64 | Low | Helical | TRUE | FALSE | FALSE |  |
| 6L9H | 2.6 | CG | I.DG-30 | J.DC30 | Low | Helical | TRUE | TRUE | TRUE |  |
| 6LE9 | 2.6 | AT | I.DA-51 | J.DT51 | Low | Helical | TRUE | FALSE | TRUE |  |
| 6LE9 | 2.6 | AT | I.DA44 | J.DT-44 | Low | Helical | TRUE | TRUE | FALSE |  |
| 6LE9 | 2.6 | AT | I.DT-5 | J.DA5 | Low | Helical | TRUE | FALSE | FALSE |  |
| 6RYI | 2.691 | AT | H.DT2 | I.DA15 | Low | Terminal-1 | TRUE | TRUE | FALSE |  |
| 6S16 | 3.409 | AT | D.DT10 | F.DA24 | Low | Terminal | TRUE | TRUE | FALSE |  |
| 6UPY | 3.4 | AT | T.DT8 | N.DA11 | Low | Helical | TRUE | TRUE | FALSE |  |
| 6W3U | 2.4 | CG | C.DG3 | E.DC9 | Low | Helical | TRUE | TRUE | TRUE |  |
| 6ZMN | 2.333 | AT | C.DA16 | D.DT1 | Low | Terminal | TRUE | TRUE | FALSE |  |
| 1F44 | 2.05 | AT | M.DA2 | N.DT19 | WC | Terminal | TRUE | TRUE | TRUE |  |
| 1NZB | 3.1 | AT | C.DT108 | D.DA129 | WC | Helical | TRUE | TRUE | FALSE |  |
| 1P51 | 2.5 | AT | E.DT16 | F.DA4 | WC | Flank | TRUE | TRUE | FALSE |  |
| 1P51 | 2.5 | AT | I.DT16 | H.DA4 | WC | Flank | FALSE | TRUE | FALSE |  |
| 1PVR | 2.65 | AT | C.DA1 | D.DT34 | WC | Terminal | TRUE | TRUE | FALSE |  |
| 1Q3U | 2.9 | AT | G.DT108 | H.DA129 | WC | Helical | TRUE | TRUE | FALSE |  |
| 1RZR | 2.8 | AT | E.DT701 | B.DA714 | WC | Terminal-1 | TRUE | FALSE | FALSE |  |
| 1SXP | 2.5 | AT | C.DT6 | D.DA21 | WC | Flank | TRUE | TRUE | FALSE |  |
| 2DY4 | 2.65 | AT | E.DA6 | F.DT112 | WC | Helical | TRUE | TRUE | FALSE |  |
| 2V9W | 3 | AT | C.DT12 | D.DA7 | WC | Pol_n-2 | TRUE | TRUE | FALSE | Weird clash of paired T |
| 2VA3 | 2.98 | AT | P.DA13 | T.DT6 | WC | Pol_n-2 | TRUE | TRUE | FALSE | Sticky_end |
| 2VJV | 1.9 | AT | C.DA25 | C.DT35 | WC | Flank-1 | TRUE | FALSE | FALSE |  |
| 2Z9O | 3.14 | AT | C.DT22 | D.DA45 | WC | Helical | TRUE | FALSE | FALSE |  |
| 3HOT | 3.25 | CG | C.DG4 | D.DC53 | WC | Terminal | TRUE | TRUE | TRUE |  |
| 3QE9 | 2.51 | AT | A.DA20 | B.DT1 | WC | Terminal | TRUE | TRUE | FALSE |  |
| 3QX3 | 2.162 | AT | C.DA1 | F.DT20 | WC | Terminal | TRUE | TRUE | TRUE |  |
| 3TS8 | 2.8 | AT | K.DT20 | L.DA33 | WC | Helical | TRUE | FALSE | FALSE |  |
| 4AIJ | 2.05 | AT | C.DA10 | D.DT13 | WC | Helical | TRUE | FALSE | FALSE |  |
| 4CRX | 2.2 | AT | C.DT20 | D.DA17 | WC | Helical | TRUE | FALSE | FALSE |  |
| 4DSJ | 2.86 | AT | C.DA1 | E.DT2 | WC | Sticky_artifact | TRUE | TRUE | FALSE |  |
| 4DSJ | 2.86 | AT | C.DT2 | E.DA1 | WC | Sticky_artifact | FALSE | TRUE | FALSE |  |
| 4G0U | 2.7 | AT | C.DA1 | F.DT20 | WC | Terminal | TRUE | TRUE | FALSE |  |
| 4J3N | 2.3 | AT | C.DA1 | F.DT20 | WC | Terminal | TRUE | TRUE | FALSE |  |
| 4MF8 | 2.32 | AT | T.DT7 | P.DA10 | WC | Pol_n-1 | TRUE | TRUE | FALSE |  |
| 4MFA | 2.27 | AT | T.DT7 | P.DA10 | WC | Pol_n-1 | TRUE | TRUE | FALSE |  |
| 4PCB | 2.5 | AT | B.DA5 | B.DT14 | WC | Helical | TRUE | FALSE | FALSE |  |
| 4XVM | 3.2 | AT | T.DA3 | P.DT11 | WC | Helical | TRUE | FALSE | FALSE |  |
| 4YPH | 2.32 | AT | B.DA2 | C.DT22 | WC | Terminal | TRUE | TRUE | FALSE |  |
| 4YPR | 2.59 | AT | C.DT22 | D.DA2 | WC | Terminal | TRUE | TRUE | FALSE |  |
| 4ZTF | 2.7 | AT | C.DA19 | D.DT1 | WC | Terminal | TRUE | TRUE | FALSE |  |
| 4ZTJ | 2.67 | AT | C.DA19 | D.DT1 | WC | Terminal | TRUE | TRUE | FALSE |  |
| 5A0M | 2.9 | AT | D.DT5 | K.DA22 | WC | Helical | TRUE | FALSE | FALSE |  |
| 5CLV | 2.5 | AT | C.DA14 | D.DT7 | WC | Helical | TRUE | FALSE | FALSE |  |
| 5DB7 | 2.209 | AT | T.DT7 | P.DA10 | WC | Pol_n-1 | TRUE | TRUE | FALSE |  |
| 5DB8 | 2.547 | AT | T.DT7 | P.DA10 | WC | Pol_n-1 | TRUE | TRUE | FALSE |  |
| 5DBB | 2.25 | AT | T.DT7 | P.DA10 | WC | Pol_n-1 | TRUE | TRUE | FALSE |  |
| 5LGY | 2.92 | AT | E.DT21 | F.DA1 | WC | Terminal | TRUE | TRUE | FALSE |  |
| 5OQO | 3.25 | AT | C.DT7 | D.DA12 | WC | Helical | TRUE | FALSE | FALSE |  |
| 5UOP | 2.85 | AT | C.DA19 | D.DT1 | WC | Terminal | TRUE | TRUE | FALSE |  |
| 5UOQ | 2.61 | AT | C.DA19 | D.DT1 | WC | Terminal | TRUE | TRUE | FALSE |  |
| 5VFX | 2.81 | AT | I.DA2 | J.DT22 | WC | Terminal-1 | TRUE | FALSE | FALSE |  |
| 5XQ2 | 3.33 | AT | E.DT7 | F.DA14 | WC | Flank | TRUE | TRUE | FALSE |  |
| 5XQ2 | 3.33 | AT | X.DT7 | Y.DA14 | WC | Flank | TRUE | TRUE | FALSE |  |
| 6DSU | 1.98 | AT | P.DT10 | T.DA4 | WC | Pol_n-1 | TRUE | TRUE | TRUE |  |
| 6FI8 | 2.598 | AT | J.DA25 | J.DT35 | WC | Flank-1 | TRUE | FALSE | FALSE |  |
| 6FQQ | 3.25 | AT | G.DA15 | H.DT2 | WC | Terminal-1 | FALSE | FALSE | FALSE |  |
| 6FQQ | 3.25 | AT | L.DA15 | M.DT2 | WC | Terminal-1 | FALSE | FALSE | FALSE |  |
| 6IFM | 2.804 | AT | M.DT9 | N.DA19 | WC | Helical | TRUE | FALSE | FALSE |  |
| 6JPI | 3.143 | AT | E.DT4 | F.DA25 | WC | Helical | TRUE | TRUE | FALSE |  |
| 6JVZ | 2.48 | AT | C.DT-2 | D.DA2 | WC | Terminal | TRUE | TRUE | FALSE |  |
| 6M3L | 2.75 | AT | C.DT-3 | C.DA3 | WC | Terminal-1 | TRUE | FALSE | FALSE |  |
| 6NCM | 2.704 | AT | C.DA11 | D.DT7 | WC | Helical | TRUE | FALSE | FALSE |  |
| 6O8G | 2.64 | AT | H.DA6 | I.DT11 | WC | Helical | TRUE | FALSE | FALSE |  |
| 6QFD | 2.133 | AT | E.DA11 | F.DT18 | WC | Helical | TRUE | FALSE | FALSE |  |
| 7CE1 | 3.2 | AT | g.DA18 | h.DT3 | WC | Terminal | TRUE | TRUE | FALSE |  |
| 7CE1 | 3.2 | AT | s.DT12 | t.DA9 | WC | Helical | TRUE | FALSE | FALSE |  |
| 7CE1 | 3.2 | AT | g.DT3 | h.DA18 | WC | Terminal | FALSE | TRUE | FALSE |  |
| 7CE1 | 3.2 | AT | o.DT3 | p.DA18 | WC | Terminal | FALSE | TRUE | FALSE |  |
| 1IC8 | 2.6 | AT | E.DA316 | F.DT406 | WC>HG | Helical | TRUE | FALSE | TRUE |  |
| 1IC8 | 2.6 | AT | E.DT306 | F.DA416 | WC>HG | Helical | TRUE | FALSE | FALSE |  |
| 1RM1 | 2.5 | AT | D.DT2 | E.DA35 | WC>HG | Terminal-1 | TRUE | FALSE | FALSE |  |
| 3CLC | 2.8 | AT | E.DT2 | F.DA34 | WC>HG | Terminal-1 | TRUE | FALSE | FALSE |  |
| 3Q05 | 2.4 | AT | K.DT10 | L.DA43 | WC>HG | Helical | FALSE | FALSE | FALSE |  |
| 3V6T | 1.85 | AT | I.DT-2 | J.DA2 | WC>HG | Terminal | FALSE | TRUE | FALSE |  |
| 4OSL | 2.447 | AT | I.DT-2 | J.DA2 | WC>HG | Terminal | FALSE | TRUE | TRUE |  |
| 4OSM | 2.454 | AT | I.DT-2 | J.DA2 | WC>HG | Terminal | FALSE | TRUE | FALSE |  |
| 4OSR | 1.944 | AT | I.DT-2 | J.DA2 | WC>HG | Terminal | FALSE | TRUE | TRUE |  |
| 4OSV | 1.996 | AT | I.DT-2 | J.DA2 | WC>HG | Terminal | FALSE | TRUE | TRUE |  |
| 4OT3 | 1.944 | AT | I.DT-2 | J.DA2 | WC>HG | Terminal | FALSE | TRUE | FALSE |  |
| 6FQQ | 3.25 | AT | G.DT16 | H.DA1 | WC>HG | Terminal | FALSE | TRUE | FALSE |  |
| 6FQQ | 3.25 | AT | L.DT16 | M.DA1 | WC>HG | Terminal | FALSE | TRUE | FALSE | Remove phosphate |
| 6JRP | 3 | AT | B.DA11 | C.DT1 | WC>HG | Terminal | FALSE | TRUE | FALSE |  |

a. This table contains a total of 241 bps = 215 bps from *Starting* + 23 augmented bps during data curation.

b. ‘HG’ = Hoogsteen; ‘HG>WC’ = ambiguous Hoogsteen; ‘HG=WC’ = ambiguous; ‘WC>HG’ = ambiguous Watson-Crick; ‘WC’ = Watson-Crick; ‘Low’ = weak density therefore ambiguous

c. ‘Terminal’ = terminal bp; ‘Terminal-1’ = the bp next to the terminal bp; ‘Helical’ = helical bp; ‘Nick’ = bp flanking a nick; ‘Nick-1’ = one bp away from a nick; ‘Pol_n-1’ = the position *n*-1 flanking the active site *n* in polymerase; ‘Pol_n-2’ = the position *n*-2 in the polymerase; ‘Flank’ = the bp flanking a DNA bulge; ‘Flank-1’ = one bp away from a DNA bulge; ‘Sticky_artifact’ = artifactual base pairing at sticky end from two nearby DNA

d. For bp entries with redundant bps, the primary entry is labeled as Primary=True while the remaining redundant entries are labeled as Primary=False

e. The bp is in chemically or structurally stressed DNA regions (see definition in main)

f. *Hintze et al.* means the bp was identified in the prior study^1^

**Supplementary Table 5. Comparison of R-factors between Watson-Crick and Hoogsteen models using TLS-based *PHENIX* refinement for all the Hoogsteen and ambiguous Hoogsteen examples identified in this study**

| PDB ID | Protein | Resolution (Å)^a^ | Space group | R-work  (Watson-Crick) | R-free  (Watson-Crick) | R-work  (Hoogsteen) | R-free  (Hoogsteen) |
| --- | --- | --- | --- | --- | --- | --- | --- |
| 1S97 | DNA polymerase IV (Dpo4) | 2.4 | P 1 21 1 | 0.1792 | 0.2463 | 0.1764 | 0.2484 |
| 2VA2 | DNA polymerase IV (Dpo4) | 2.8 | P 1 21 1 | 0.1648 | 0.2370 | 0.1611 | 0.2335 |
| 2ZHG | Redox-sensitive transcriptional activator soxR | 2.8 | P 61 2 2 | 0.2197 | 0.2963 | 0.2191 | 0.2968 |
| 3JXB | Repressor protein C2 | 1.67 | P 21 21 21 | 0.1795 | 0.2157 | 0.1757 | 0.2115 |
| 3Q05 | Cellular tumor antigen p53 | 2.4 | P 21 21 2 | 0.1964 | 0.2414 | 0.1939 | 0.2396 |
| 3QQY | Ribosomal protein 3/homing endonuclease-like protein fusion | 2.401 | P 21 21 21 | 0.1743 | 0.2259 | 0.1733 | 0.2291 |
| 3V6H | DNA polymerase IV (Dpo4) | 2.3 | P 1 21 1 | 0.1977 | 0.2546 | 0.1956 | 0.2586 |
| 3V6J | DNA polymerase IV (Dpo4) | 2.3 | P 1 21 1 | 0.2009 | 0.2605 | 0.1980 | 0.2555 |
| 3V6T | dHax3 | 1.85 | P 1 21 1 | 0.1838 | 0.2200 | 0.1829 | 0.2218 |
| 4G82 | Tumor protein p73 | 3.1 | P 61 | 0.1768 | 0.2333 | 0.1704 | 0.2298 |
| 4GC7 | DNA polymerase IV (Dpo4) | 2.89 | P 1 21 1 | 0.1789 | 0.2687 | 0.1783 | 0.2740 |
| 4GJR | dHax3 | 1.85 | P 1 21 1 | 0.1847 | 0.2294 | 0.1840 | 0.2305 |
| 4IWR | Regulatory protein Esp1396I | 2.4 | P 32 | 0.1910 | 0.2685 | 0.1875 | 0.2622 |
| 4KPY | TtAgo | 2.406 | P 21 21 21 | 0.1864 | 0.2373 | 0.1838 | 0.2365 |
| 4NCA | TtAgo | 2.489 | P 21 21 21 | 0.1834 | 0.2427 | 0.1806 | 0.2410 |
| 4NCB | TtAgo | 2.189 | P 1 21 1 | 0.1717 | 0.2349 | 0.1704 | 0.2355 |
| 4OSL | dHax3 | 2.447 | P 1 21 1 | 0.1859 | 0.2546 | 0.1844 | 0.2591 |
| 4OSM | dHax3 | 2.454 | P 1 21 1 | 0.1817 | 0.2491 | 0.1821 | 0.2484 |
| 4OSR | dHax3 | 1.944 | P 1 21 1 | 0.1826 | 0.2284 | 0.1826 | 0.2299 |
| 4OSV | dHax3 | 1.996 | P 1 21 1 | 0.1874 | 0.2334 | 0.1870 | 0.2346 |
| 4OT3 | dHax3 | 1.944 | P 1 21 1 | 0.1799 | 0.2216 | 0.1792 | 0.2220 |
| 5A0W | Homing endonuclease I-DMOI | 2.2 | P 1 21 1 | 0.1552 | 0.1947 | 0.1549 | 0.1976 |
| 5E6C | Glucocorticoid receptor | 2.2 | P 21 21 21 | 0.2103 | 0.2497 | 0.2063 | 0.2473 |
| 5EYO | Protein max | 2.39 | C 1 2 1 | 0.2020 | 0.2344 | 0.2057 | 0.2406 |
| 5HP4 | T5 flap Endonuclease | 1.86 | P 43 21 2 | 0.1768 | 0.2028 | 0.1751 | 0.2007 |
| 5WN0 | Human APE1 | 2.6 | P 1 | 0.1930 | 0.2781 | 0.1910 | 0.2772 |
| 5ZDZ | HMGB1 A-B box/mouse RAG1/mouse RAG2 | 2.8 | P 1 21 1 | 0.1750 | 0.2305 | 0.1719 | 0.2283 |
| 6FQP | Homeobox protein TGIF1 | 2.42 | P 21 21 21 | 0.2131 | 0.2539 | 0.2154 | 0.2566 |
| 6FQQ | Homeobox protein TGIF1 | 3.25 | P 21 21 21 | 0.2058 | 0.2642 | 0.2046 | 0.2754 |
| 6JRP | Protein capicua homolog | 3 | P 32 | 0.2530 | 0.3377 | 0.2493 | 0.3394 |

a. The resolutions are obtained directly from the RCSB PDB.

**Supplementary Table 6. Non-redundant Hoogsteen base pairs identified in this study**

| PDB ID | Bp | nt_1 | nt_2 | Context | Stress | WC  clash | C1′-C1′  WC (Å) | C1′-C1′  HG (Å) | Note |
| --- | --- | --- | --- | --- | --- | --- | --- | --- | --- |
| 1S97 | AT | E.DT12 | I.DA7 | Pol_n-2 | TRUE | TRUE | 8.692 | 8.285 | Hintze *et al.* |
| 3JXB | CG | A.DC1 | B.DG40 | Terminal | TRUE | TRUE | 8.666 | 8.577 | Hintze *et al.* |
| 3QQY | CG | B.DC25 | C.DG1 | Terminal | TRUE | TRUE | 9.823 | 8.662 |  |
| 3V6H | AT | C.DA7 | D.DT12 | Pol_n-2 | TRUE | TRUE | 8.92 | 8.531 |  |
| 3V6J | AT | P.DT12 | B.DA7 | Pol_n-2 | TRUE | TRUE | 8.83 | 8.519 | Hintze *et al.* |
| 3V6T | AT | G.DT-2 | H.DA2 | Terminal | TRUE | TRUE | 8.794 | 8.031 | Hintze *et al.* |
| 4GJR | AT | G.DT1 | H.DA15 | Terminal | TRUE | TRUE | 9.268 | 8.086 |  |
| 4IWR | AT | C.DA1 | D.DT25 | Terminal | TRUE | TRUE | 9.116 | 8.7 |  |
| 4KPY | AT | C.DT14 | N.DA6 | Helical | FALSE | FALSE | 9.843 | 9.148 |  |
| 4OSL | AT | G.DT-2 | H.DA2 | Terminal | TRUE | TRUE | 9.306 | 8.615 |  |
| 4OSM | AT | G.DT-2 | H.DA2 | Terminal | TRUE | TRUE | 9.359 | 8.642 |  |
| 4OSR | AT | G.DT-2 | H.DA2 | Terminal | TRUE | TRUE | 8.763 | 8.169 |  |
| 4OT3 | AT | G.DT-2 | H.DA2 | Terminal | TRUE | TRUE | 9.234 | 8.286 |  |
| 5A0W | CG | B.DG11 | C.DC15 | Helical | TRUE | TRUE | 8.847 | 7.966 |  |
| 5HP4 | AT | X.DT5 | X.DA6 | Helical | TRUE | TRUE | 8.188 | 8.132 |  |
| 5WN0 | AT | C.DT1 | E.DA11 | Nick | TRUE | TRUE | 8.335 | 8.039 |  |
| 5WN0 | CG | C.DC2 | E.DG10 | Nick-1 | TRUE | TRUE | 9.557 | 8.146 |  |

**Supplementary Table 7. Non-redundant ambiguous Hoogsteen base pairs identified in this study**

| PDB ID | Bp | nt_1 | nt_2 | Context | Stress | WC  clash | C1′-C1′  WC (Å) | C1′-C1′  HG (Å) | Note |
| --- | --- | --- | --- | --- | --- | --- | --- | --- | --- |
| 2VA2 | AT | C.DT12 | D.DA7 | Pol_n-2 | TRUE | TRUE | 9.184 | 8.905 |  |
| 2ZHG | AT | B.DT10 | B.DA11 | Helical | FALSE | TRUE | 9.528 | 9.276 |  |
| 3Q05 | AT | K.DA19 | L.DT34 | Helical | FALSE | TRUE | 9.204 | 8.736 |  |
| 3Q05 | AT | K.DT20 | L.DA33 | Helical | FALSE | FALSE | 9.501 | 8.988 |  |
| 3V6H | AT | C.DA8 | D.DT11 | Helical | FALSE | FALSE | 10.115 | 9.893 |  |
| 3V6J | AT | P.DT11 | B.DA8 | Helical | FALSE | TRUE | 9.954 | 9.741 |  |
| 4G82 | AT | E.DA404 | F.DT415 | Helical | FALSE | TRUE | 9.447 | 9.049 |  |
| 4GC7 | CG | E.DC13 | F.DG6 | Pol_n-2 | FALSE | TRUE | 9.005 | 8.842 |  |
| 4GC7 | CG | E.DC14 | F.DG5 | Pol_n-1 | TRUE | TRUE | 9.999 | 9.771 |  |
| 4NCA | AT | C.DT14 | G.DA3 | Helical | FALSE | TRUE | 9.282 | 8.737 |  |
| 4NCB | AT | C.DT14 | D.DA6 | Terminal-1 | TRUE | TRUE | 8.304 | 7.933 |  |
| 4OSV | AT | G.DT-2 | H.DA2 | Terminal | TRUE | FALSE | 9.305 | 8.488 |  |
| 5E6C | AT | C.DA1 | D.DT16 | Terminal | TRUE | TRUE | 9.368 | 8.723 |  |
| 5EYO | AT | B.DA4 | B.DT19 | Terminal | TRUE | FALSE | 9.65 | 9.056 |  |
| 5ZDZ | AT | F.DA32 | I.DT15 | Nick-1 | TRUE | TRUE | 9.059 | 8.832 |  |
| 5ZDZ | AT | J.DT15 | G.DA43 | Nick-1 | TRUE | TRUE | 9.665 | 8.799 |  |
| 6FQP | AT | L.DA15 | M.DT2 | Terminal-1 | FALSE | TRUE | 9.133 | 8.703 |  |
| 6FQP | AT | L.DT16 | M.DA1 | Terminal | TRUE | FALSE | 9.624 | 8.755 |  |
| 6FQQ | AT | G.DA1 | H.DT16 | Terminal | TRUE | TRUE | 8.109 | 8.084 |  |
| 6FQQ | AT | G.DT2 | H.DA15 | Terminal-1 | TRUE | TRUE | 8.487 | 8.156 |  |
| 6JRP | AT | H.DA11 | I.DT1 | Terminal | TRUE | TRUE | 9.862 | 9.965 |  |

**Supplementary Table 8. PDB ID of Watson-Crick containing Dpo4 structures without lesions**

1JX4

1N48

1S0N

1S9F

2AGQ

2ATL

3PR4

3PR5

4F4W

4F4Z

4QW8

5YUR

5YUS

5YUT

5YUU

5YUV

5YUW

5YUX

5YUY

5YUZ

5YV0

5YV1

5YV2

5YV3

5ZLV

6IG1

**Supplementary Table 9. DNA global shape for structures with Hoogsteen base pairs and their corresponding Watson-Crick structures**

| PDB  ID | Protein | Type | nt_1 | nt_2 | RMSD  1  (Å) | RMSD  2  (Å) | α_h_  (°) | β_h_  (°) | γ_h_  (°) | ζ_h_  (°) | Minor Groove  (Å) | Major Groove  (Å) |
| --- | --- | --- | --- | --- | --- | --- | --- | --- | --- | --- | --- | --- |
| 5GQ9 | TtAgo | WC | C.DT14 | D.DA6 | 0.9 | 0.62 | 12 | 30.5 | 17.6 | 29.6 |  | 15.552 |
| 5GQ9 |  |  | E.DT14 | F.DA6 | 0.89 | 0.81 | -0.3 | 35.9 | 28.5 | 28.2 |  | 15.447 |
| 4NCB |  | HG | E.DT14 | H.DA6 | 0.92 | 0.63 | 2.3 | 33.6 | 24.6 | 26.9 |  | 15.042 |
| 4NCA |  |  | C.DT14 | G.DA3 | 0.92 | 0.6 | 6.9 | 33.2 | 22.3 | 29.2 |  |  |
| 4NCA |  |  | E.DT14 | H.DA6 | 0.89 | 0.63 | 7 | 31.6 | 21.8 | 28.8 |  |  |
| 4KPY |  |  | C.DT14 | N.DA6 | 0.88 | 0.72 | 0 | 35.5 | 28.5 | 28.5 |  |  |
| 4KPY |  |  | E.DT14 | M.DA6 | 0.93 | 0.67 | 5.1 | 34.7 | 23.3 | 28.4 |  | 15.329 |
| 4UN9 | I-DMOI | WC | B.DT12 | C.DA14 | 0.96 | 0.84 | -178.3 | 29.3 | -133.7 | 48 | 5.444 | 19.78 |
| 4UN9 |  |  | E.DT12 | F.DA14 | 0.99 | 0.95 | 179.1 | 28.5 | -129 | 50.1 | 5.444 | 21.199 |
| 4UN9 |  |  | H.DT12 | I.DA14 | 0.98 | 0.8 | -178.7 | 28.7 | -132.2 | 49.1 | 5.398 | 19.557 |
| 5A0W |  | HG | B.DT12 | C.DA14 | 1.41 | 0.99 | 142.6 | 43.4 | -104 | 38.6 | 4.858 | 23.685 |
| 5A0W |  |  | E.DT12 | F.DA14 | 1.42 | 0.89 | 149 | 41.7 | -110.4 | 38.6 | 4.908 | 23.245 |
| 5A0W |  |  | H.DT12 | I.DA14 | 1.43 | 0.98 | 145.5 | 42.4 | -107.2 | 38.3 | 4.901 | 23.527 |
| 5WN1 | Human APE1 | WC | --- | E.DT12 | 0.83 | 0.66 | 45.6 | 36.7 | -18.8 | 26.8 | 15.086 | 21.373 |
| 5WN4 |  |  | D.C7R10 | E.DT12 | 0.8 | 0.66 | 40.3 | 33.3 | -10.4 | 29.9 | 14.634 | 20.195 |
| 5WN0 |  | HG | D.DC10 | E.DG12 | 0.7 | 1.55 | 21.4 | 36.9 | -3.7 | 17.7 | 17.229 | 20.86 |
| 5HP4 | T5 flap Endonuclease | HG | X.DT5 | X.DA6 | 0.94 | 1.33 | -28.2 | 15 | 65.3 | 37.1 | 11.503 | 17.798 |
| 2ZHG | soxR | HG | B.DT10 | B.DA11 | 1.24 | 1.14 | -6.2 | 60.3 | 27.3 | 21.1 | 17.772 | 16.745 |

**Supplementary Table 10. Spin lock power and offsets used in the *R*_1ρ_ experiments**

| **Nuclei** | **[spin lock power] {offset frequencies}** |
| --- | --- |
|  | **[ω_SL_ 2π^-1^(s^-1^)] {Ω 2π^-1^(s^-1^)}** |
| **hpCG** | |
| G6-C8 | [150, 200, 250, 300, 350, 400, 500, 600, 700, 800, 900, 1000, 1200, 1400, 1600, 1800, 2000, 2500, 3000, 3500] {0}; [150] {-451, -410, -369, -328, -287, -246, -205, -164, -123, -82, -41, -10, 10, 41, 82, 123, 164, 205, 246, 287, 328, 369, 410, 451}; [300] {-902, -820, -738, -656, -574, -492, -410, -328, -246, -164, -82, -10, 10, 82, 164, 246, 328, 410, 492, 574, 656, 738, 820, 902}; [500] {-1496, -1360, -1224, -1088, -952, -816, -680, -544, -408, -272, -136, -10, 10, 136, 272, 408, 544, 680, 816, 952, 1088, 1224, 1360, 1496}; [900] {-1496, -1360, -1224, -1088, -952, -816, -680, -544, -408, -272, -136, -10, 10, 136, 272, 408, 544, 680, 816, 952, 1088, 1224, 1360, 1496} |
| **hpTA** | |
| G6-C8 | [100] {-352, -320, -288, -256, -224, -192, -160, -128, -96, -64, -32, -10, 10, 32, 64, 96, 128, 160, 192, 224, 256, 288, 320, 352}; [200] {-704, -640, -576, -512, -448, -384, -320, -256, -192, -128, -64, -10, 10, 64, 128, 192, 256, 320, 384, 448, 512, 576, 640, 704}; [300] {-1045, -950, -855, -760, -665, -570, -475, -380, -285, -190, -95, -10, 10, 95, 190, 285, 380, 475, 570, 665, 760, 855, 950, 1045}; [500] {-1749, -1590, -1431, -1272, -1113, -954, -795, -636, -477, -318, -159, -10, 10, 159, 318, 477, 636, 795, 954, 1113, 1272, 1431, 1590, 1749}; [700] {-1749, -1590, -1431, -1272, -1113, -954, -795, -636, -477, -318, -159, -10, 10, 159, 318, 477, 636, 795, 954, 1113, 1272, 1431, 1590, 1749} |
| **hpTG** | |
| G6-C8 | [100] {-352, -320, -288, -256, -224, -192, -160, -128, -96, -64, -32, -10, 10, 32, 64, 96, 128, 160, 192, 224, 256, 288, 320, 352}; [200] {-704, -640, -576, -512, -448, -384, -320, -256, -192, -128, -64, -10, 10, 64, 128, 192, 256, 320, 384, 448, 512, 576, 640, 704}; [300] {-1045, -950, -855, -760, -665, -570, -475, -380, -285, -190, -95, -10, 10, 95, 190, 285, 380, 475, 570, 665, 760, 855, 950, 1045}; [500] {-1749, -1590, -1431, -1272, -1113, -954, -795, -636, -477, -318, -159, -10, 10, 159, 318, 477, 636, 795, 954, 1113, 1272, 1431, 1590, 1749}; [700] {-1749, -1590, -1431, -1272, -1113, -954, -795, -636, -477, -318, -159, -10, 10, 159, 318, 477, 636, 795, 954, 1113, 1272, 1431, 1590, 1749} |
| **hpTT** | |
| G6-C8 | [100] {-352, -320, -288, -256, -224, -192, -160, -128, -96, -64, -32, -10, 10, 32, 64, 96, 128, 160, 192, 224, 256, 288, 320, 352}; [200] {-704, -640, -576, -512, -448, -384, -320, -256, -192, -128, -64, -10, 10, 64, 128, 192, 256, 320, 384, 448, 512, 576, 640, 704}; [300] {-1045, -950, -855, -760, -665, -570, -475, -380, -285, -190, -95, -10, 10, 95, 190, 285, 380, 475, 570, 665, 760, 855, 950, 1045}; [500] {-1749, -1590, -1431, -1272, -1113, -954, -795, -636, -477, -318, -159, -10, 10, 159, 318, 477, 636, 795, 954, 1113, 1272, 1431, 1590, 1749}; [700] {-1749, -1590, -1431, -1272, -1113, -954, -795, -636, -477, -318, -159, -10, 10, 159, 318, 477, 636, 795, 954, 1113, 1272, 1431, 1590, 1749} |

**Supplementary Table 11. Exchange parameters obtained from 2-state fitting of the off-resonance ^13^C *R*_1ρ_ relaxation dispersion NMR data**

| Construct | Nuclei | Δω_B_ (ppm) | *p*_B_ (%) | *k*_ex_ (s^-1^) | *R*_1_ (s^-1^) | *R*_2_ (s^-1^) | Red. χ^2^ |
| --- | --- | --- | --- | --- | --- | --- | --- |
| hpCG | G6-C8 | 3.38±0.05 | 0.467±0.042 | 668±74 | 4.25±0.03 | 17.94±0.04 | 0.75 |
| hpTA | G6-C8 | 3.04±0.02 | 0.477±0.042 | 556±69 | 3.17±0.01 | 17.89±0.05 | 2.49 |
| hpTG | G6-C8 | 2.97±0.03 | 1.449±0.072 | 803±65 | 2.70±0.07 | 14.92±0.24 | 0.49 |
| hpTT | G6-C8 | 3.22±0.02 | 4.634±0.073 | 1225±39 | 3.45±0.06 | 18.93±0.34 | 0.92 |

**Supplementary Table 12. Van der Waals contacts to the purine base with the Hoogsteen versus Watson-Crick model for polymerase Dpo4 detected by DNAproDB^14^.**

| **HG model** | | | | | | |
| --- | --- | --- | --- | --- | --- | --- |
| PDB ID | distance | nt | nt_atom | nt_moiety | aa | aa_atom |
| 1S97 | 3.87 | I.DA7 | C2 | majorgw | A.ILE248 | CG2 |
|  | 3.869 | J.DA7 | C2 | majorgw | B.ILE248 | CG2 |
| 3V6H | 3.825 | C.DA8 | C2 | majorgw | A.ARG336 | NE |
|  | 3.8 | C.DA8 | C2 | majorgw | A.ARG336 | NH2 |
|  | 3.802 | C.DA8 | C2 | majorgw | A.ARG336 | CZ |
|  | 3.798 | C.DA8 | N3 | majorgw | A.ARG336 | NH2 |
|  | 3.66 | C.DA8 | N3 | majorgw | A.ARG336 | CZ |
|  | 3.899 | C.DA8 | N3 | majorgw | A.ARG336 | NH1 |
| 3V6J | 3.895 | B.DA8 | C2 | majorgw | A.ARG336 | NH1 |
|  | 3.864 | B.DA8 | N3 | majorgw | A.ARG336 | NH1 |
|  | 3.851 | B.DA8 | N3 | majorgw | A.ARG336 | NE |
|  | 3.598 | B.DA8 | N3 | majorgw | A.ARG336 | CZ |
|  | 3.742 | B.DA8 | N3 | majorgw | A.ARG336 | NH2 |
| 4GC7 | 3.273 | F.DG6 | N2 | majorgw | B.ILE248 | CG2 |
|  | 3.691 | F.DG5 | C2 | majorgw | B.ARG332 | NH2 |
|  | 3.031 | F.DG5 | N2 | majorgw | B.ARG332 | NH1 |
|  | 3.185 | F.DG5 | N2 | majorgw | B.ARG332 | CZ |
|  | 2.928 | F.DG5 | N2 | majorgw | B.ARG332 | NH2 |
|  | 3.641 | F.DG5 | N3 | majorgw | B.ARG332 | NH2 |
|  | 3.355 | D.DG6 | N2 | majorgw | A.ILE248 | CG2 |
|  | 3.703 | D.DG5 | C2 | majorgw | A.ARG332 | NH2 |
|  | 3.035 | D.DG5 | N2 | majorgw | A.ARG332 | NH2 |
|  | 3.217 | D.DG5 | N2 | majorgw | A.ARG332 | CZ |
|  | 2.912 | D.DG5 | N2 | majorgw | A.ARG332 | NH1 |
|  | 3.581 | D.DG5 | N3 | majorgw | A.ARG332 | NH2 |
| **WC model** | | | | | | |
| PDB ID | distance | nt | nt_atom | nt_moiety | aa | aa_atom |
| 3V6H | 3.702 | C.DA8 | C8 | majorgw | A.ARG336 | NE |
|  | 3.4 | C.DA8 | C8 | majorgw | A.ARG336 | CZ |
|  | 3.183 | C.DA8 | C8 | majorgw | A.ARG336 | NH2 |
|  | 3.663 | T.DA8 | C8 | majorgw | B.ARG336 | CZ |
|  | 3.64 | T.DA8 | C8 | majorgw | B.ARG336 | NH2 |
| 3V6J | 3.808 | B.DA8 | C8 | majorgw | A.ARG336 | NE |
|  | 3.626 | B.DA8 | C8 | majorgw | A.ARG336 | CZ |
|  | 3.672 | B.DA8 | C8 | majorgw | A.ARG336 | NH2 |
|  | 3.723 | M.DA8 | C8 | majorgw | J.ARG336 | CZ |
|  | 3.825 | M.DA8 | C8 | majorgw | J.ARG336 | NH1 |
| 4GC7 | 3.296 | F.DG5 | C8 | majorgw | B.ARG332 | NH2 |
|  | 3.664 | F.DG5 | N7 | majorgw | B.ARG332 | NH2 |
|  | 3.4 | D.DG5 | C8 | majorgw | A.ARG332 | NH2 |
